# Distinct NMDA Receptor Pools Determine Diverse Forms of Cortical Plasticity

**DOI:** 10.64898/2026.02.14.705915

**Authors:** Sabine Rannio, Kalenga Lubembele, Claire Bokang Ko, Nicole Cherepacha, Vivian Yuwei Li, Gemma Moffat, Shawniya Alageswaran, Aurore Thomazeau, Rafael Luján, P. Jesper Sjöström

## Abstract

NMDA receptors (NMDARs) are well-established as coincidence detectors in Hebbian learning, but this requires postsynaptic localization. Presynaptic NMDARs, however, remain controversial due to limitations of pharmacology and calcium imaging. We therefore dissected the function of distinct NMDAR pools using more direct techniques, including immunogold EM, sparse genetic deletion, and paired recordings from layer-5 (L5) pyramidal cell (PC) synapses in mouse primary visual cortex (V1). We found that pre- but not postsynaptic NMDARs regulate both spontaneous and evoked neurotransmitter release. In spike-timing-dependent plasticity, we uncovered a double dissociation: timing-dependent long-term depression (tLTD) requires pre- but not postsynaptic NMDARs, whereas timing-dependent long-term potentiation (tLTP) needs post- but not presynaptic NMDARs. Postsynaptic NMDAR loss also caused developmentally delayed dendritic spine loss and altered axonal and dendritic architecture. In summary, NMDARs do not act as a unified plasticity signal but rather confer location-specific control over diverse forms of synaptic signaling, plasticity, and circuit structure.

## INTRODUCTION

Memory formation in the brain and information processing by neural circuits are shaped by experience through diverse forms of neuronal plasticity, which operate across multiple timescales and affect distinct facets of neuronal function.^1–3^

On timescales of minutes to days, neurons remodel their axonal and dendritic branches, and dendritic spines are formed or eliminated — processes collectively referred to as structural plasticity.^4,5^ Changes in synaptic strength that persist over similar timescales are known as long-term synaptic plasticity,^1,2^ which is often closely associated with structural modifications of dendritic spines,^6,7^ linking functional and anatomical circuit refinement. A well-studied variant is spike-timing-dependent plasticity (STDP), in which the temporal order of pre- and postsynaptic activity determines whether synaptic strength increases or decreases.^8,9^ In neocortical layer-5 (L5) pyramidal cell (PC) pairs, pre-before-post firing induces timing-dependent LTP (tLTP), whereas the reverse order yields timing-dependent LTD (tLTD).^10,11^

On sub-second to multi-second timescales, synapses also express short-term plasticity driven by presynaptic spiking and release dynamics.^3,12^ Even in the absence of spiking, synapses release vesicles spontaneously, a form of transmission that is also plastic.^13^

Many forms of synaptic and circuit plasticity rely on NMDARs. Because they require both glutamate binding and postsynaptic depolarization to open,^14,15^ NMDARs detect coincident pre- and postsynaptic activity — a core feature of Hebbian learning.^16^ But for Hebbian coincidence detection, NMDARs must be positioned postsynaptically, where they sense presynaptic glutamate release and postsynaptic depolarization.^17–19^ Yet evidence over the past three decades has shown that there are also presynaptic NMDARs.^20–24^

Several studies have proposed that presynaptic NMDARs can impact both long- and short-term plasticity across multiple brain regions.^18,25,26^ However, alternative interpretations have challenged this view, suggesting that NMDARs attributed to presynaptic terminals may instead reside in dendrites of presynaptic cells,^27^ be postsynaptic,^28^ or may not exist.^29^ Their precise role and even their existence thus remain debated.^18,25,26^

This controversy may stem from several unusual properties of NMDARs. Firstly, NMDARs frequently contain subunits or operate under conditions that limit Ca^2+^ permeability,^30^ reducing the detectability of their activation. Moreover, both postsynaptic and presynaptic NMDARs can engage in non-ionotropic signaling,^31–38^ which produces no detectable Ca^2+^ transients and which is insensitive to channel-pore blockers such as MK-801 as well as to Ca^2+^ chelators such as BAPTA. Pharmacological specificity can also be challenging. For example, intracellular MK-801 incompletely blocks NMDAR currents.^39^ When coupled with low Ca^2+^ permeability,^30^ Ca^2+^ imaging and pharmacology thus become blunt instruments for assigning plasticity to specific pre- versus postsynaptic NMDAR pools.

To overcome these limitations, we relied on more direct techniques, including immuno-electron microscopy (EM) and whole-cell recordings of connected L5 PC → PC pairs combined with sparse genetic NMDAR deletion. This approach allowed us to systematically link distinct pre- and postsynaptic NMDAR pools to specific forms of plasticity in V1 L5 PCs, including synaptic release, short-term dynamics, tLTD, and tLTP. We also assessed structural changes in axodendritic arborizations and spine numbers. Together, our findings provide direct loss-of-function evidence that helps settle a long-standing controversy and show that NMDAR signaling is not monolithic but follows a compartment-specific logic that supports diverse forms of cortical plasticity.

## RESULTS

### Direct ultrastructural evidence for pre- and postsynaptic NMDARs

Previous studies, including our own, have provided pharmacological and Ca^2+^ imaging evidence for presynaptic NMDARs in L5 PC axons,^18,25,26^ although other work has challenged their existence.^29^ We reasoned that immuno-EM of excitatory synapses in V1 L5 could help resolve this discrepancy by providing more direct evidence,^20,22,40,41^ while simultaneously revealing distribution patterns of NMDAR subunits on either side of the synaptic cleft.

We therefore carried out post-embedding immunogold labeling of GluN1, GluN2A and GluN2B subunits in ultrathin sections of postnatal day (P) 21 mice (see Methods). This confirmed the presence of all three subunits pre- as well as postsynaptically at excitatory synapses in V1 L5 (Figure 1A, B). Across sections, we measured the distance of immunogold particles from the postsynaptic density and observed an apparent asymmetry in the relative distribution of GluN2A and GluN2B (Figure 1B). To quantify this asymmetry, we used a Monte Carlo approach to test whether the observed spatial distributions differed from those expected by chance (Figure 1C). This analysis revealed a greater relative presynaptic preponderance of GluN2B over GluN2A (Figure 1C), even though both subunits were overall enriched postsynaptically (Figure 1B).

**Figure 1.**
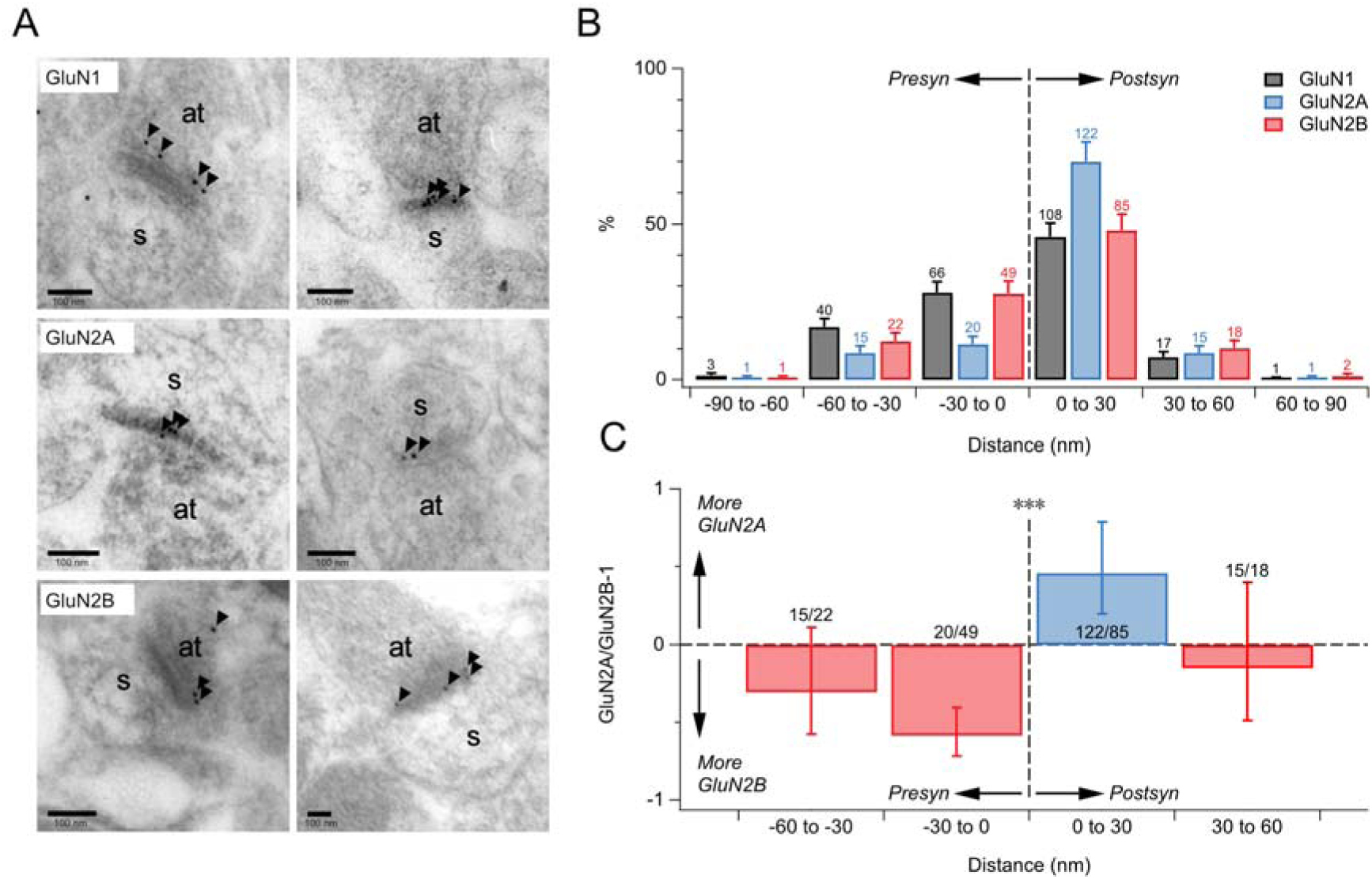
NMDARs are found both pre- and postsynaptically at L5 excitatory synapses. (A) Post-embedding immunogold EM of V1 L5 sections from P21 mice showed GluN1, GluN2A, and GluN2B subunits in presynaptic axon terminals (at) and postsynaptic spines (s) (arrowheads). Synapses were identified ultrastructurally and quantified if immunogold-labeled (see Methods). (B) Quantification of immunogold particle locations from morphologically identified excitatory synapses, pooled across three mice, revealed that GluN1 was distributed similarly across pre- and postsynaptic compartments (postsynaptic: 126/235 immunoparticles = 54%, χ^2^ p = 0.27, n = 102 excitatory synapses). GluN2A was enriched postsynaptically (138/174 = 79%, χ^2^ p < 0.001, n = 90), whereas GluN2B showed a more modest postsynaptic bias (105/177 = 59%, χ^2^ p < 0.05, n = 90). Error bars: normalized square root of counts. (C) GluN2B was ∼2× more likely than GluN2A to be found presynaptically (72 vs. 36 puncta; p < 0.001; Monte Carlo randomization of particle identities across pooled locations; see Methods). Puncta outside ± 60 nm are not shown. Error bars were computed by error propagating from (B).

Together, these data demonstrate that NMDARs are present on both sides of V1 L5 excitatory synapses within V1 layer 5, with GluN2B- and GluN2A-containing receptors relatively enriched pre- and postsynaptically, respectively. Although these synapses were sampled broadly from L5, the presynaptic enrichment of GluN2B is consistent with GluN2B-selective pharmacology acting presynaptically at V1 L5 PC → PC synapses at this age.^37,38,42–44^

### Regulation of spontaneous release by pre- but not postsynaptic NMDARs

Having established that NMDARs reside on both sides of excitatory synapses in V1 L5, we next asked how these distinct receptor pools contribute to functional forms of plasticity. We began with the regulation of spontaneous release, which — based on pharmacology — has long been attributed to presynaptic NMDARs.^21,24,42^ However, this interpretation has been debated,^27,29^ and the role of postsynaptic NMDARs in spontaneous release regulation has remained unclear.

To test the presynaptic contribution directly, we established the triple-transgenic mouse line Emx1^cre/+^;NR1^flox/flox^;Ai9^tdTom/+^ (Supplementary Figure S1) which as expected^38,45^ deleted NMDARs relatively globally, meaning from >90% of excitatory neurons (denoted Del, Supplementary Figures S2-3). We then compared spontaneous release in PCs from Del slices and in PCs from WT slices, which were obtained from Emx^cre/+^;NR1^+/+^;Ai9^tdTom/+^ littermates.

In PCs from WT littermates, wash-in of the NMDAR antagonist AP5 reversibly decreased mEPSC frequency but not amplitude (Figure 2), consistent with presynaptic NMDARs enhancing spontaneous release, in agreement with our prior studies.^38,42,43^ In contrast, AP5 did not reduce mEPSC frequency in PCs from Del mice (Figure 2), suggesting a need for presynaptic NMDARs. However, because postsynaptic NMDARs were likely also removed in this mouse model, a potential postsynaptic contribution could not be excluded.

**Figure 2.**
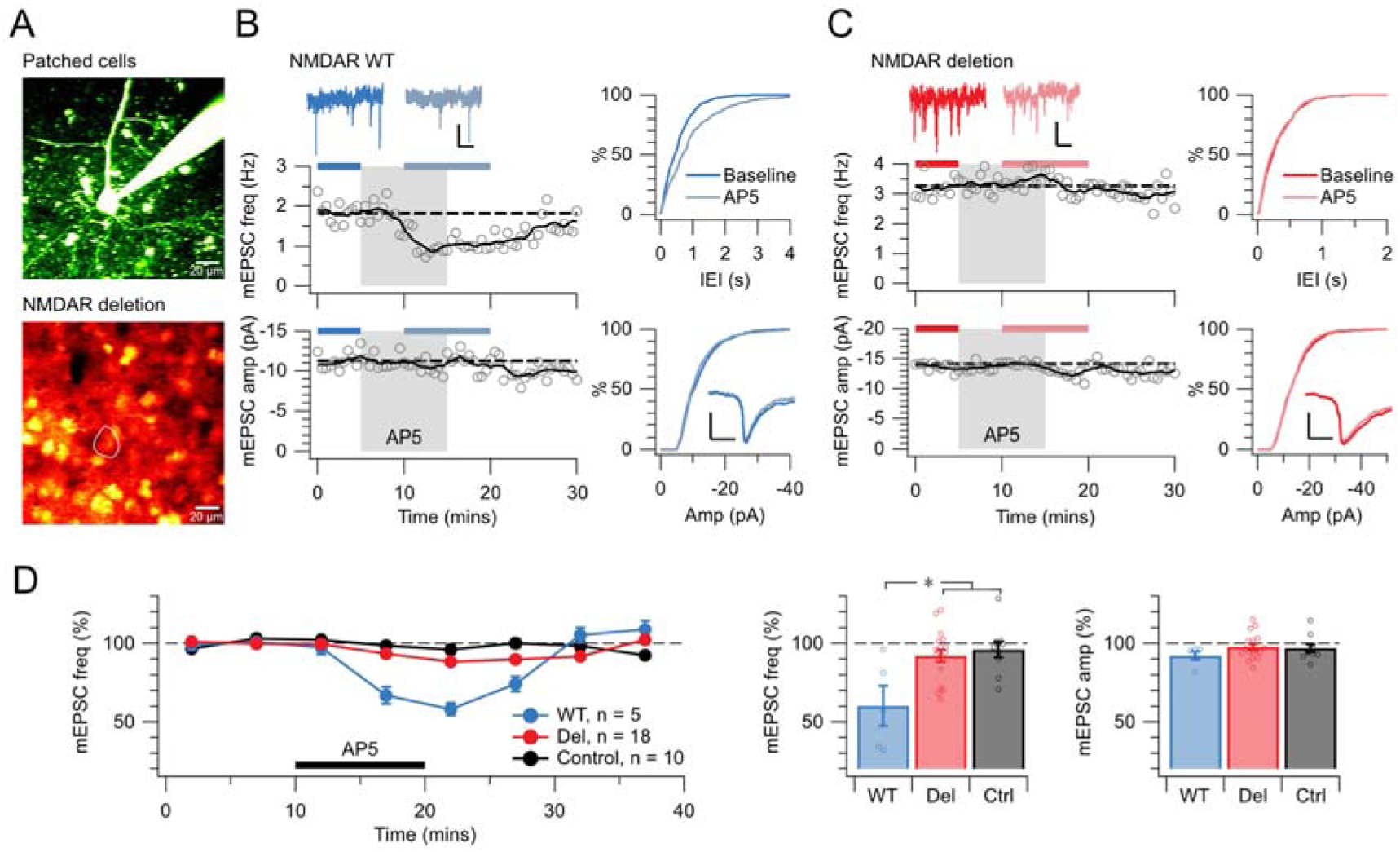
Global NMDAR deletion suggests presynaptic control of spontaneous release. (A) In this sample whole-cell recording (top) from a triple-transgenic Emx1^Cre/+^;NR1^flox/flox^;Ai9^tdTomato/+^ mice (bottom), NMDARs were globally deleted, which we denote ‘Del’ below. Recordings obtained from Emx1^Cre/+^;NR1^+/+^;Ai9^tdTomato/+^ littermates (images not shown) are denoted ‘WT’. (B) In this representative PC recording from a WT L5 mouse, AP5 reversibly reduced mEPSC frequency (top, KS test p < 0.001) but not amplitude (bottom, p = 0.09), consistent with reduced presynaptic release. Sample traces scale bars: 200 ms, 10 pA. Insets show mEPSC traces averaged over periods indicated in blue and light blue. Scale bars: 5 ms, 5 pA. (C) For the sample PC in (A) obtained from a Del mouse, AP5 had no effect on mEPSC frequency (top, KS test p = 0.82) or amplitude (bottom, p = 0.9), suggesting that the effect of AP5 in WT requires presynaptic NMDARs. Scale bars as in (B). (D) AP5 robustly reduced mEPSC frequency in PCs from WT mice relative to no-drug controls (ANOVA p < 0.01), but had no detectable effect in PCs from Del mice (p = 0.52), consistent with presynaptic NMDAR control of spontaneous release. Baseline mEPSC frequencies were indistinguishable (WT: 2.7 ± 0.2 Hz, n = 62; Del: 2.8 ± 0.1 Hz, n = 86; Mann Whitney p = 0.40), and mEPSC amplitudes were unaffected.

We therefore turned to the GluN2B-selective NMDAR antagonist Ro 25-6981, which blocks pre- but not postsynaptic NMDARs at juvenile V1 L5 PC → PC synapses^37,38,42–44^ and which aligns with our EM evidence for greater presynaptic GluN2B component (Figure 1). The GluN2B-selective antagonist Ro 25-6981 similarly reduced mEPSC frequency, but not amplitude, in WT PCs but had no effect in Del PCs (Supplementary Figure S4). These findings are consistent with a presynaptic GluN2B-containing NMDAR contribution to the regulation of spontaneous release.

To directly test postsynaptic NMDAR involvement, we developed a sparse NMDAR deletion model. Injection of AAV9-eSYN-mCherry-T2A-iCre-WPRE or AAV9-eSYN-EGFP-T2A-iCre-WPRE virus into V1 of NR1^flox/flox^ neonates (see Methods) reliably removed NMDARs in a sparse subset of PCs by postnatal day (P) 10 (Supplementary Figures S5-6). This approach allowed us to record from NMDAR-deleted PCs while leaving the vast majority of presynaptic PCs unaffected (Supplementary Figures S5-6). Comparing mEPSCs in NMDAR-deleted PCs to neighboring unlabeled WT PCs revealed no differences in frequency or amplitude (Figure 3A–E). Thus, postsynaptic NMDARs did not contribute to the regulation of spontaneous vesicle fusion.

**Figure 3.**
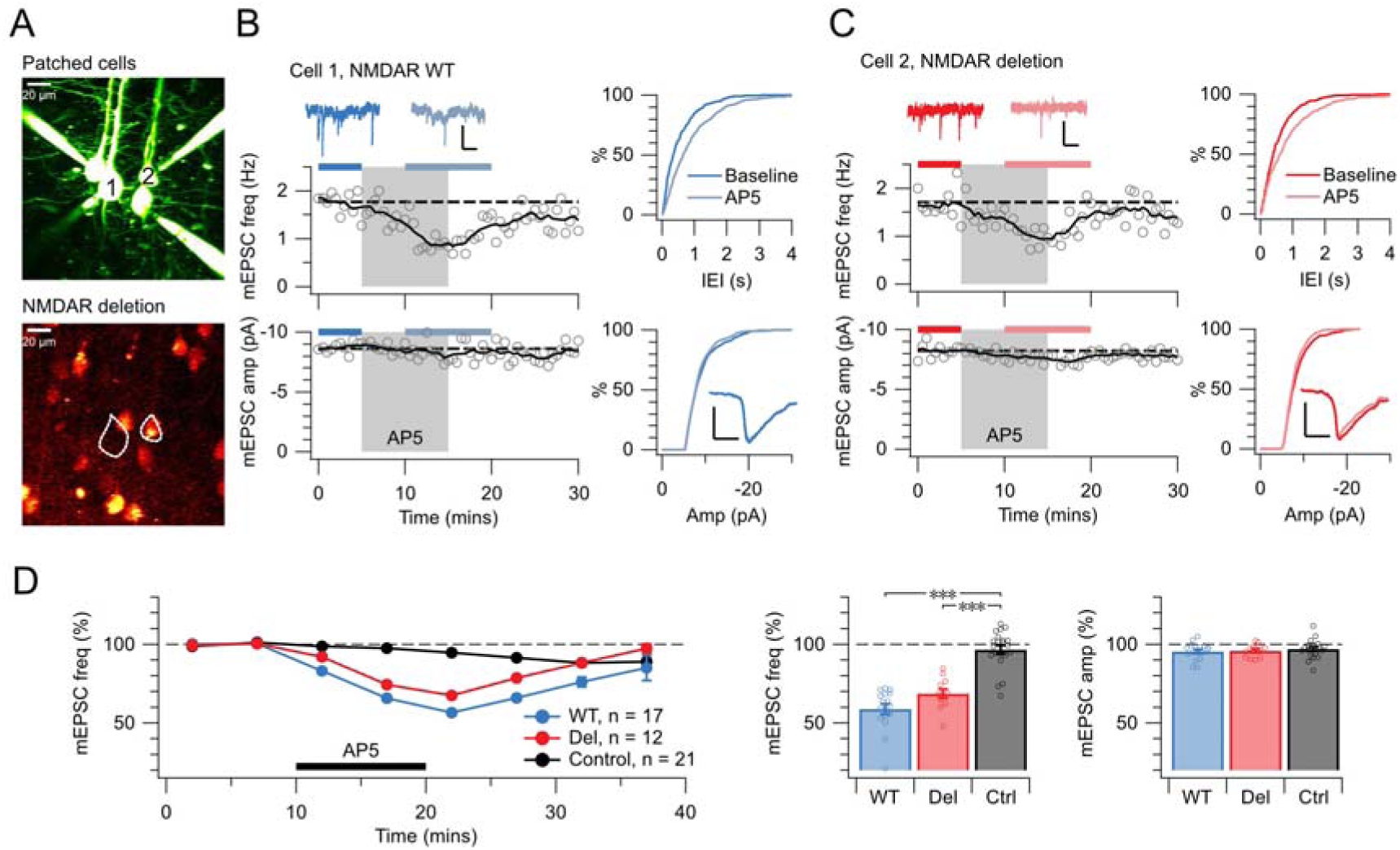
Sparse NMDAR deletion reveals no postsynaptic role in spontaneous release. (A) In this sample quadruple recording (top), PC1 but not PC2 had intact NMDARs following sparse viral deletion in NR1^flox/flox^ mice (see Methods). Fluorescent labelling allowed identification of NMDAR-deleted neurons (bottom). (B) In WT PC1, AP5 reversibly reduced mEPSC frequency (top, KS test p < 0.001) without affecting amplitude (bottom, p = 0.15), indicating a presynaptic reduction in release. Sample traces scale bars: 200 ms, 10 pA. Insets show averaged mEPSC traces. Scale bars: 5 ms, 5 pA. (C) In the paired NMDAR-deleted PC2, AP5 similarly reduced frequency (top, KS test p < 0.001) but not amplitude (bottom, p = 0.15). With postsynaptic NMDARs deleted, the remaining AP5 effect excludes a postsynaptic role. Scale bars as in (B) (D) AP5 consistently reduced frequency in WT and NMDAR-deleted PCs relative to controls (ANOVA p < 0.001), demonstrating that NMDARs controlling spontaneous release were not postsynaptic. Baseline mEPSC frequencies were indistinguishable (WT: 2.9 ± 0.2 Hz, n = 63; Del: 2.9 ± 0.3 Hz, n = 38, Mann Whitney p = 0.62), and amplitudes were unaffected.

Together, these complementary deletion strategies demonstrate that spontaneous release is regulated by pre- but not postsynaptic NMDARs. By selectively removing postsynaptic receptors while preserving most presynaptic NMDARs, our approach excludes any contribution of postsynaptic NMDAR signaling — whether ionotropic or non-ionotropic^18,25,26^ — to the control of spontaneous release.

### Evoked release is controlled by presynaptic, but not by postsynaptic, NMDARs

We next investigated how presynaptic and postsynaptic NMDAR pools contribute to evoked glutamate release. Using our sparse genetic deletion strategy, we recorded from synaptically connected L5 PC → PC pairs in which NMDARs were selectively removed from the presynaptic neuron (Del → WT), the postsynaptic neuron (WT → Del), or left intact in both neurons (WT → WT; Figure 4). Pairs are thus labeled by presynaptic → postsynaptic genotype.

**Figure 4.**
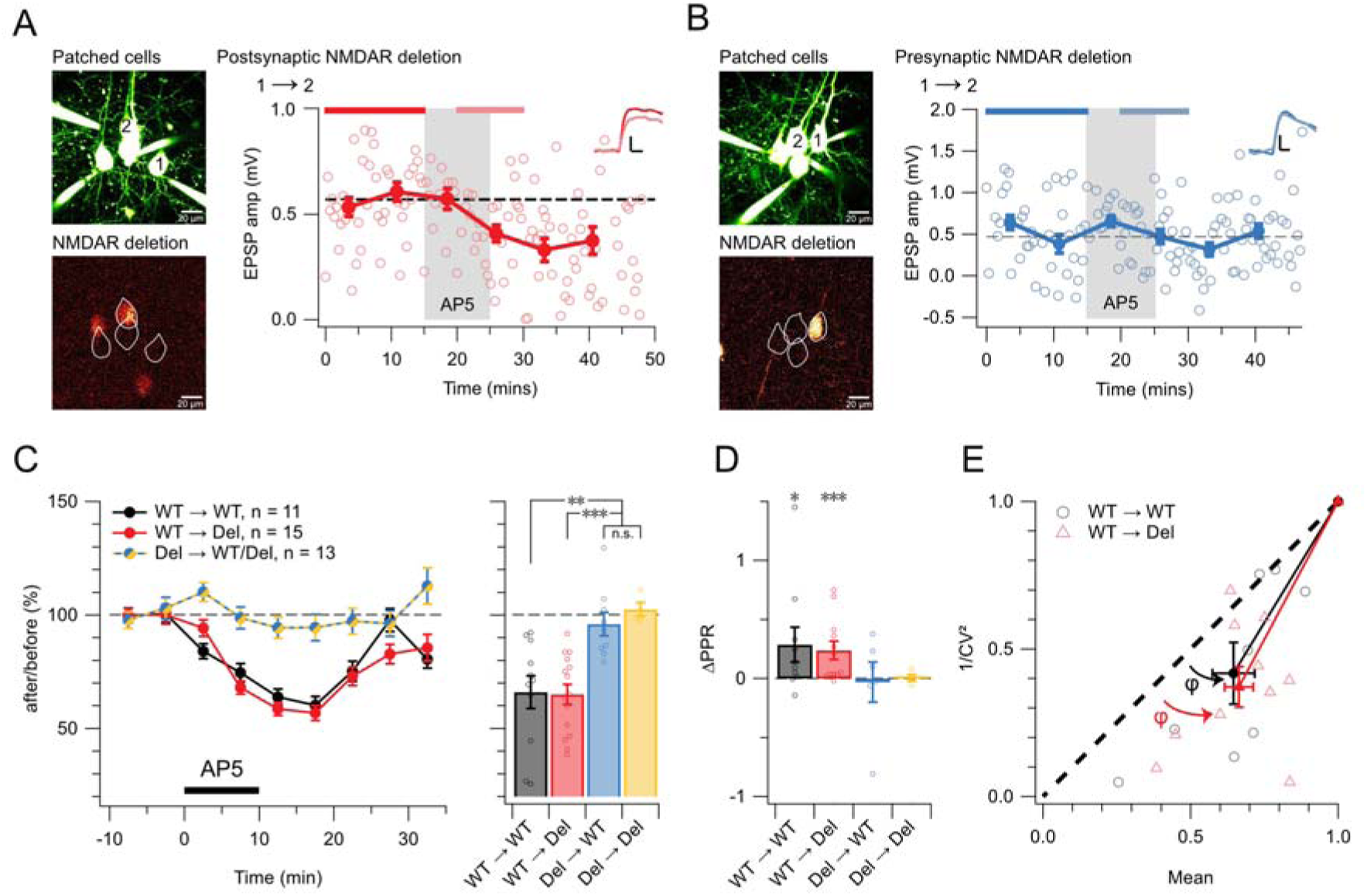
Pre- but not postsynaptic NMDARs regulate evoked release. (A) Sample PC1 → PC2 pair with postsynaptic NMDAR deletion identified by fluorescent tag. AP5 wash-in reduced EPSP amplitude (light red: 0.43 ± 0.04 mV vs red: 0.57 ± 0.03 mV, Student’s t p < 0.01), indicating that postsynaptic NMDARs were not required for the AP5 effect. Inset scale bars: 5 ms, 0.2 mV. (B) Sample PC1 → PC2 pair with presynaptic NMDAR deletion identified by fluorescent tag, in which AP5 had no effect on EPSP amplitude (light blue: 0.48 ± 0.07 mV vs blue: 0.51 ± 0.08 mV, p = 0.76). Scale bars as in (A). (C) Across the population, AP5 wash-in reduced EPSP amplitude in WT → Del pairs and WT → WT control pairs, but not in Del → WT/Del pairs (ANOVA, p < 0.001). Category plots show mean; whiskers indicate SEM. (D) PPR increased following AP5 wash-in in WT → Del pairs and WT → WT controls but was unchanged in Del → WT pairs or Del → Del pairs, consistent with a presynaptic locus of action. Connections with baseline EPSP amplitudes < 0.3 mV were excluded. (E) CV analysis further supported a presynaptic locus of AP5-induced EPSP reduction in WT → Del pairs (red: φ = 16 ± 4°, Wilcoxon p < 0.01) and WT → WT controls (black: φ = 13 ± 4°, p < 0.05), because, for synaptic weakening, points below the diagonal imply reduced release probability.^46^ Del → WT and Del → Del pairs are omitted as AP5 had no effect.

Blocking NMDARs with AP5 reduced EPSP amplitude in WT → WT and WT → Del pairs (Figure 4A, C), consistent with a presynaptic reduction in release probability.^37,38,42,43^ In contrast, AP5 had no effect on EPSP amplitude in Del → WT pairs (Figure 4B, C), indicating that removal of presynaptic but not postsynaptic NMDARs abolished the sensitivity of evoked transmission to AP5.

Paired-pulse measurements further supported a presynaptic locus: AP5 increased the paired-pulse ratio (PPR) in WT → WT and WT → Del pairs, consistent with a decrease in release probability, whereas PPR was unchanged in Del → WT pairs (Figure 4D). Coefficient-of-variation (CV) analysis^46^ likewise pointed toward a presynaptic change in release (Figure 4E).

Together, these results demonstrate that presynaptic, but not postsynaptic, NMDARs regulate evoked glutamate release at L5 PC → PC synapses, in agreement with our earlier studies.^37,38,42,43^

### Pre- but not postsynaptic NMDARs are required for tLTD

Post-before-pre spiking, with postsynaptic spikes preceding presynaptic spikes by 10-25 milliseconds, induces tLTD at L5 PC → PC pairs.^10^ To determine whether pre- or postsynaptic NMDARs were required for tLTD, we induced tLTD in slices with sparse NMDAR deletion.

We found that WT → WT and WT → Del pairs reliably expressed tLTD, as evidenced by a sustained reduction in EPSP amplitude following induction (Figure 5A, C). Consistent with a presynaptic locus of expression, CV analysis indicated a reduction in release probability following tLTD induction (Figure 5D). In contrast, Del → WT and Del → Del pairs failed to express tLTD, with post-induction EPSP amplitudes indistinguishable from baseline (Figure 5B, C).

**Figure 5.**
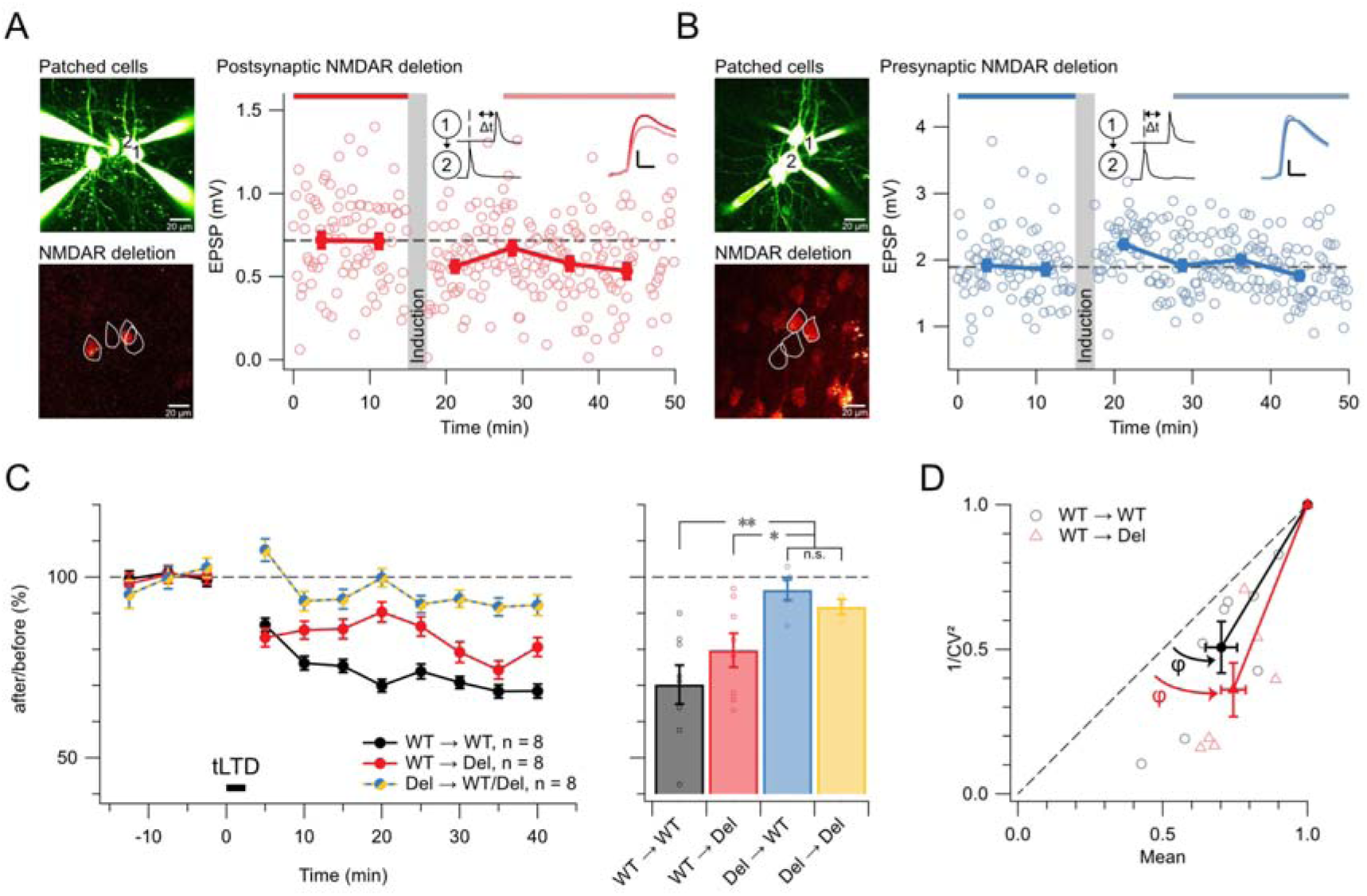
Pre- but not postsynaptic NMDARs are required for tLTD. (A) For this sample PC1 → PC2 recording (top left) with postsynaptic NMDAR deletion (bottom left), 1-Hz induction with a Δt = -25 ms spike-timing difference yielded tLTD (light red: 0.6 ± 0.03 mV vs red: 0.7 ± 0.03 mV, Student’s *t* test p < 0.01). Inset scale bars: 5 ms, 0.2 mV. (B) In contrast, in this representative PC1 → PC2 pair with presynaptic deletion, EPSP amplitude was unchanged by the same induction protocol (light blue: 1.9 ± 0.04 mV vs blue: 1.9 ± 0.05 mV, Student’s *t* test p = 0.90). Average EPSP traces inset top right highlight absence of plasticity. Inset scale bars: 5 ms, 0.5 mV. (C) Sparse genetic deletion of presynaptic NMDARs in PC → PC pairs reliably abolished tLTD compared to WT → WT pairs (ANOVA p < 0.01), while postsynaptic NMDAR deletion pairs showed robust tLTD (80 ± 5%). (D) CV analysis was consistent with a presynaptic locus of tLTD expression for both WT → WT pairs (φ = 13 ± 3°, Wilcoxon p < 0.01) and WT → Del pairs (φ = 22 ± 3°, Wilcoxon p < 0.05)

Together, these results demonstrate that tLTD at L5 PC → PC synapses depends on presynaptic but *not* on postsynaptic NMDARs, thereby extending and reaffirming our prior findings highlighting a need for presynaptic NMDARs.^42,47^

### Post- but not presynaptic NMDARs are required for tLTP

We next examined the contribution of distinct NMDAR pools to tLTP, which is classically thought to require postsynaptic NMDAR activation, whereas a role for presynaptic NMDARs has remained unclear.^8^

Using a high-frequency pre-before-post induction protocol, we reliably induced tLTP in Del → WT pairs and in control WT → WT pairs, but not in WT → Del pairs (Figure 6A–C). Thus, deletion of postsynaptic but not presynaptic NMDARs abolished tLTP. As reported previously, weaker synapses potentiated more than stronger ones (Supplementary Figure S7), consistent with saturation of potentiation at high initial synaptic strength.^10,48–53^ Analysis of CV indicated a mixed locus of tLTP expression (Figure 6D,E), consistent with prior findings in rat V1.^54^ CV analysis of Del → WT pairs suggested a presynaptic shift, but this observation did not meet criteria for nonparametric tests and is reported descriptively.

**Figure 6.**
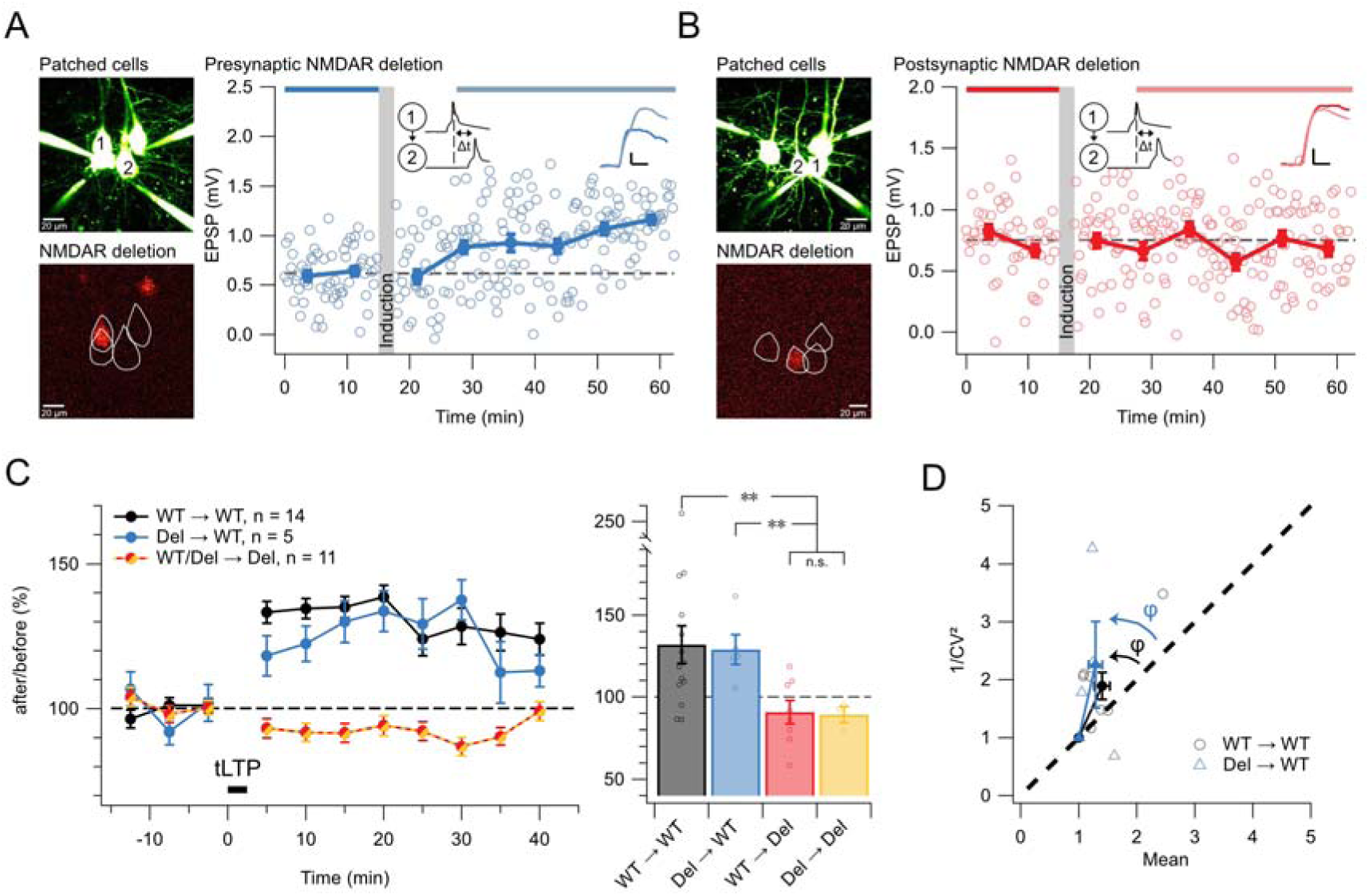
Post- but not presynaptic NMDARs are required for tLTP. (A) Sample Del → WT pair, with presynaptic NMDAR deletion indicated by fluorescent labelling (bottom left), showed robust tLTP following 50-Hz pre-before-post induction (Δt = +10 ms; light blue: 1.0 ± 0.03 mV vs blue: 0.6 ± 0.03 mV, Student’s t p < 0.001). Inset scale bars: 5 ms, 0.2 mV. (B) In contrast, in a sample WT → Del pair, the same induction protocol failed to elicit tLTP (light red: 0.70 ± 0.03 mV vs red: 0.75 ± 0.04 mV, Student’s t p = 0.38). (C) Across the population, WT → Del pairs did not potentiate (Student’s t p = 0. 24 versus 100%) nor did Del → Del pairs (p = 0.16 versus 100%), whereas Del → WT and control WT → WT pairs did (Welch ANOVA p < 0.01). Potentiation in Del → WT pairs was indistinguishable from WT → WT controls (Student’s t p = 0.84). (D) CV analysis in WT → WT pairs did not deviate from the diagonal (φ = 17 ± 7°, Wilcoxon p = 0.08), consistent with mixed pre- and postsynaptic expression of tLTP.^46,54^ Nevertheless, 1/CV^2^ increased (1.9 ± 0.2, Wilcoxon p < 0.01), suggesting a robust presynaptic component of potentiation.^55^ CV analysis in Del → WT pairs suggested mixed expression (φ: 11 ± 28°, Wilcoxon p = 0.88; normalized 1/CV^2^: 2.3 ± 0.8, Wilcoxon p = 0.25) but was not statistically conclusive (n = 4).

These findings reveal a division of labor in which post- but not presynaptic NMDARs underpin tLTP induction, whereas expression involves both pre- and postsynaptic changes. Together with the reciprocal requirement for presynaptic NMDARs in tLTD, this establishes a double dissociation in STDP, with distinct NMDAR-based coincidence detectors for tLTP and tLTD.

### NMDARs regulate basal dendritic spine density in L5 pyramidal neurons

Because long-term plasticity is associated with the formation and stabilization of dendritic spines at excitatory synapses,^4–6^ we asked whether the disruption of NMDAR signaling — a key driver of synaptic plasticity^1,2,30^ — was accompanied by changes in dendritic spine density.

In the global NMDAR deletion model, basal dendritic spine density was reduced in L5 PCs lacking NMDARs compared to WT controls of juvenile animals aged P12-17 (Figure 7A, B). These findings suggest that NMDAR signaling contributes to the developmental accumulation or maintenance of basal spines on L5 PCs.

**Figure 7.**
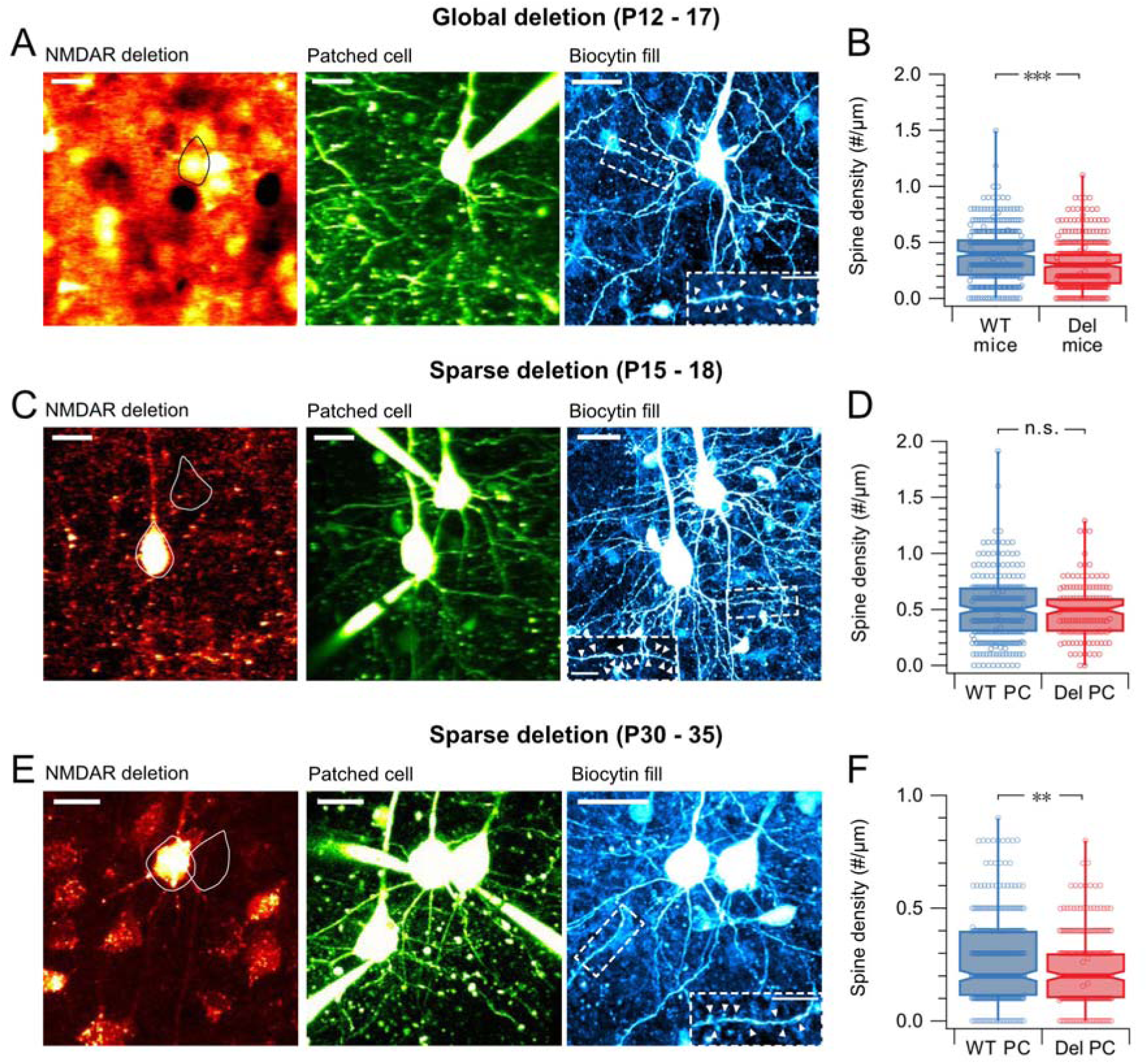
NMDAR deletion reduces basal dendritic spine density in L5 PCs. (A) Experimental approach for spine quantification in globally NMDAR-deleted and WT L5 PCs. Left: tdTomato fluorescence identified excitatory neurons. Middle: live 2-photon imaging of Alexa 488 during whole-cell recording and biocytin filling. Right: post hoc confocal imaging following histological processing with Alexa 488-conjugated streptavidin, resolving dendritic spines (arrowheads). (B) Global NMDAR deletion reduced basal spine density in juvenile mice compared to WT littermates (Mann-Whitney p < 0.001, WT: n = 9 cells, 7 slices, 5 animals; Del: n = 6 animals, 10 slices, 14 cells). Each data point represents a 10-μm basal dendritic segment. Box plots show median and interquartile range with whiskers indicating min/max values. (C) Experimental approach for spine analysis following sparse NMDAR deletion. Left: mCherry fluorescence identified NMDAR-deleted PCs. Middle: live 2-photon imaging of Alexa 488 during whole-cell recording and biocytin filling. Right: post hoc confocal imaging after histological processing with Alexa-647-conjugated streptavidin resolved dendritic spines (arrowheads). (D) Sparse NMDAR deletion did not reduce basal spine density in juvenile mice (Mann-Whitney p = 0.58, n = 13 cells, 8 slices, 6 animals) (E) As in (C), spine analysis in adolescent mice following sparse postsynaptic NMDAR deletion, with histological processing (right) carried out with Alexa-488-conjugated streptavidin. (F) Sparse NMDAR deletion reduced basal spine density in adolescent mice (Mann-Whitney p < 0.01, n = 13 cells, 8 slices, 4 animals).

However, because global NMDAR deletion eliminates receptors throughout the local network, this approach could not distinguish pre- from postsynaptic contributions. We therefore next examined dendritic spine density following sparse, postsynaptic NMDAR deletion. In juvenile animals (P15-18), sparse postsynaptic NMDAR deletion surprisingly did not lower overall basal dendritic spine density (Figure 7C, D). Notably, tLTP was already abolished at this stage in WT → Del pairs (Figure 6), indicating that the plasticity deficit precedes and is not a consequence of structural remodeling.

To independently assess the lack of an early spine-density phenotype, we statistically analyzed all our paired recordings (Figures 4-6), which revealed no detectable difference in connectivity across WT → WT and WT → Del pairs (Supplementary Figure S8). Likewise, baseline spontaneous release rate — a proxy for the number of inputs onto a neuron — was not affected by global or sparse postsynaptic NMDAR deletion (Table S1). These observations lent support to our spine density analysis after sparse postsynaptic NMDAR deletion (Figure 7C, D).

We reasoned that this spine density discrepancy in global versus sparse deletion might reflect differences in the timing of NMDAR loss. Emx1-driven recombination begins embryonically at ∼E10.5,^45^ whereas sparse deletion by neonatal viral injection begins at P0–P2 (see Methods). This ∼10–12-day difference could permit early spine development to proceed relatively normally before receptor loss in the sparse deletion condition.

To test this possibility, we examined spine density in adolescent animals (P30–35) following sparse postsynaptic NMDAR deletion. At this later stage, basal dendritic spine density was robustly reduced in NMDAR-deleted PCs relative to WT controls (Figure 7E, F). A parsimonious interpretation is that postsynaptic NMDAR signaling is required for the long-term establishment or maintenance of basal dendritic spines, with deficits emerging gradually after receptor loss.

Together, our findings demonstrated that postsynaptic NMDARs contributed to the formation and stabilization of excitatory connectivity in PCs. Connectivity rate analyses additionally suggest that presynaptic NMDARs influence connection probability (Supplementary Figure S8).

### NMDARs differentially determine axonal and dendritic branching

The integration of neurons into cortical circuits depends not only on synaptic strength and connectivity, but also on the geometry of axonal and dendritic arbors. The principal cortical output neurons, L5 PCs, elaborate complex morphologies that support both intracortical and interlaminar communication.^56,57^ Because diminished NMDAR activation has been linked to reduced axonal and dendritic branching during development,^58,59^ we revisited this question with sparse cell-specific NMDAR deletion and quantitative reconstruction of L5 PC morphologies.

We first analyzed sparse NMDAR deletion in juvenile mice (Figure 8), and then tested the robustness of these effects across deletion strategies (global deletion; Supplementary Figure S9) and developmental stage (adolescent mice; Supplementary Figure S10).

**Figure 8.**
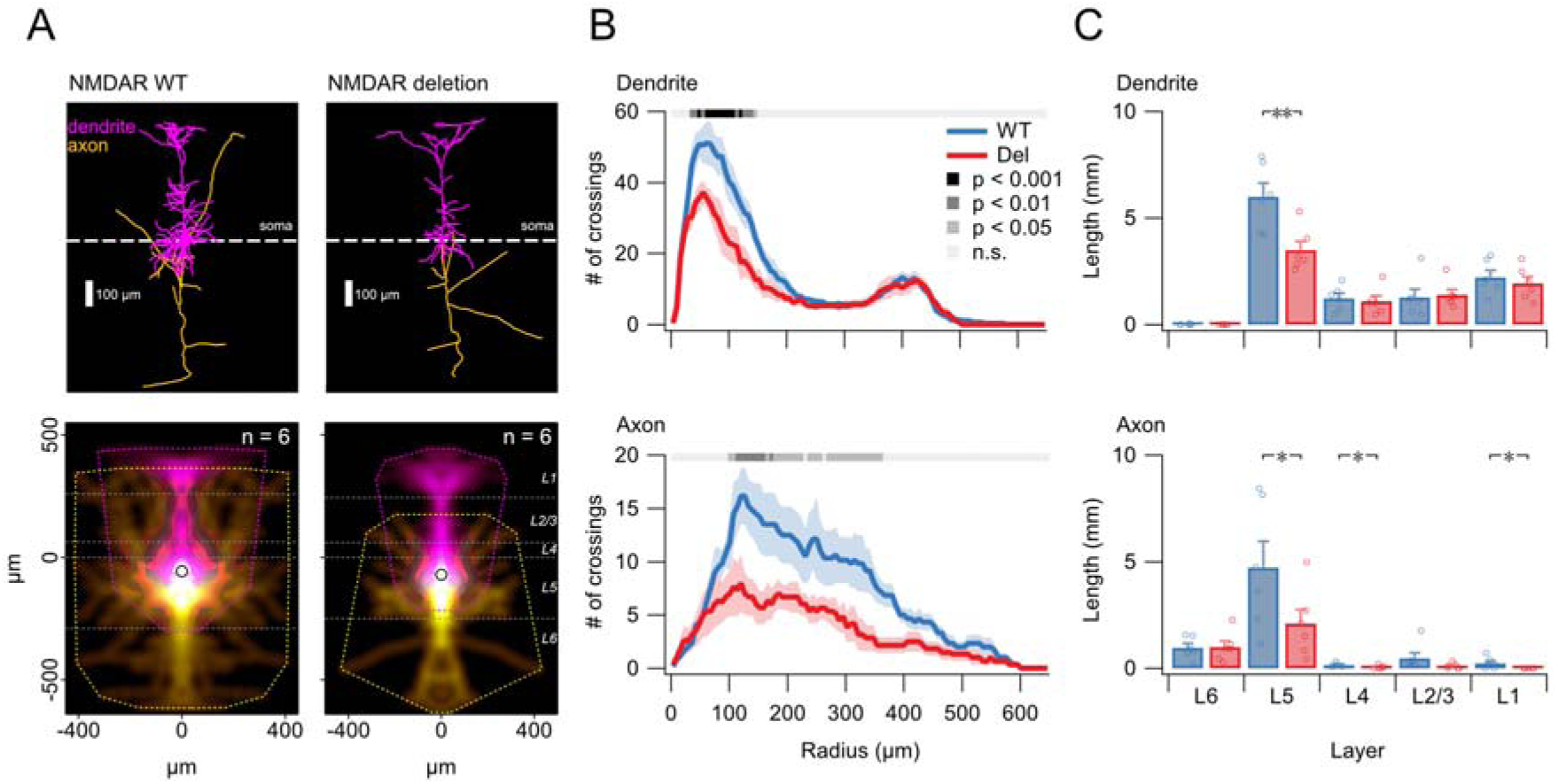
NMDARs distinctly determine axonal and dendritic branching. (A) Sample morphologies of a WT PC (left) and a PC with sparse genetic NMDAR deletion in juvenile mice, aligned on somata (dashed line). Bottom: Density maps show average extent of axonal (yellow) and dendritic (magenta) arborizations, while the convex hulls (yellow/magenta dotted lines) denote their maximum extents. Horizontal grey dotted lines demarcate neocortical layer boundaries. Open circles denote mean soma position. (B) In juvenile mice, linear mixed-effects model (LMM) analysis of Sholl plot profiles demonstrated that NMDAR deletion specifically disrupted proximal dendritic branching (top) as well as axonal branching (bottom). LMM p values indicated by grey-to-black line. (C) Sparse NMDAR deletion reduced dendritic branching specifically in L5 (top), whereas axonal arborizations were reduced across multiple layers, including L1, L4 and L5.

Using 3D reconstructions of sparsely NMDAR-deleted L5 PCs and in-slice WT controls, Sholl analysis revealed that NMDAR deletion reduced axonal branching and proximal dendritic complexity, while leaving the overall dendritic arbor extent largely intact (Figure 8). Layer-resolved analyses showed reduced axonal branching in L1, L2/3, and L5, whereas dendritic branching was selectively altered within L5.

Consistent with these findings, global NMDAR deletion in juvenile mice likewise reduced proximal dendritic branching, with differences most prominent in L5 (Supplementary Figure S9). Morphometric analyses in adolescent animals yielded comparable patterns following sparse NMDAR deletion, indicating that these effects persist across developmental stages (Supplementary Figure S10).

Together, these results demonstrate that NMDARs differentially shape axonal and dendritic arbors of L5 PCs in a layer-specific manner. NMDAR deletion preferentially reduced axonal branching across layers, whereas dendritic branching was selectively disrupted within L5. Because these anatomical effects did not track our measured changes in release or STDP (Figures 2–6), they likely reflect longer-timescale roles of NMDARs in synapse stabilization and circuit maintenance.

## DISCUSSION

In this study, we dissected the distinct contributions of pre- and postsynaptic NMDARs to multiple forms of cortical plasticity. By combining ultrastructural localization with sparse genetic deletion and paired electrophysiological recordings, we performed loss-of-function tests that pinpointed which pre- versus postsynaptic receptor population is required for spontaneous and evoked release, STDP, and longer-term structural organization. Together, our findings show that, in juvenile and adolescent V1, NMDAR signaling is not monolithic. Instead, NMDARs follow a compartment-specific logic whereby pre- and postsynaptic receptors support distinct forms of synaptic transmission, plasticity, and circuit organization.

### A division of labor between pre- and postsynaptic NMDARs

Using ultrastructural localization together with compartment-specific genetic deletion, we demonstrate that presynaptic NMDARs regulate neurotransmitter release and presynaptic plasticity, whereas postsynaptic NMDARs are dispensable for release but required for associative plasticity. Presynaptic NMDARs regulated both spontaneous neurotransmitter release and evoked release probability at L5 PC → PC synapses. These findings extend early pharmacological evidence implicating presynaptic NMDARs in release regulation,^21,24,42^ while directly excluding a contribution from postsynaptic receptors. In contrast, selective removal of postsynaptic NMDARs left spontaneous and evoked release intact.

In STDP, we found that the roles of presynaptic and postsynaptic NMDARs diverged from classical single-detector models of STDP.^8,9^ Presynaptic NMDARs were required for tLTD, consistent with a presynaptic expression of synaptic weakening,^42,47^ whereas postsynaptic NMDARs were not. By contrast, tLTP required postsynaptic NMDARs for induction, while its expression involved coordinated pre- and postsynaptic changes, consistent with prior work in neocortex and hippocampus.^11,54,60,61^ Consistent with the well-established coupling between NMDAR-dependent plasticity and structural remodeling, postsynaptic NMDAR loss led to delayed reductions in basal dendritic spine density and altered axonal and dendritic branching.^4,6,7^

By directly separating pre- and postsynaptic NMDAR pools, our study helps explain long-standing ambiguity about their respective roles in cortical circuits. Together, our findings establish a compartment-specific logic for NMDAR function, in which presynaptic receptors shape neurotransmitter release and synaptic weakening, while postsynaptic receptors support associative strengthening and longer-term stabilization of circuit structure.^2,16,62^

### Revisiting the role of presynaptic NMDARs in cortical synapses

The existence and functional relevance of presynaptic NMDARs at cortical synapses have been debated for decades. Early pharmacological and physiological studies reported NMDAR-dependent modulation of neurotransmitter release and short-term plasticity, consistent with a presynaptic locus of action at excitatory synapses.^20,21,23,24^ Subsequent work, however, challenged these interpretations, arguing that such effects could arise from dendritic or postsynaptic NMDARs, glutamate spillover, or indirect network effects.^27–29^ As a result, presynaptic NMDARs have remained controversial, with their presence and function often viewed as method- or context-dependent.^18,25,26^

A major source of this controversy lies in the limitations of commonly used experimental approaches. Calcium imaging relies on detecting local Ca^2+^ influx, yet many NMDARs — particularly those containing GluN2B or GluN3 subunits, or operating under depolarization-limited conditions — exhibit weak Ca^2+^ permeability despite being functionally engaged.^30^ Pharmacological tools are similarly constrained: channel pore blockers such as MK-801 incompletely block NMDAR signaling when applied intracellularly and do not interfere with non-ionotropic signaling modes,^31,39^ while competitive antagonists such as AP5 act globally and cannot distinguish pre- from postsynaptic receptor pools in recurrent cortical circuits. Together, these factors complicate compartment-specific interpretation of NMDAR-dependent effects.

Our study overcomes these limitations by combining ultrastructural localization with compartment-specific genetic deletion and direct functional assays. Immuno-EM provided anatomical evidence for NMDAR subunits on both sides of the synaptic cleft, consistent with ultrastructural demonstrations of pre- and postsynaptic NMDARs at hippocampal mossy fiber synapses.^41^ Sparse genetic deletion further enabled selective removal of NMDARs from either presynaptic or postsynaptic neurons without perturbing the surrounding network. Coupling these approaches with measurements of synaptic transmission and plasticity allowed causal assignment of function to each compartment.

Together, these findings reconcile prior discrepancies in the literature by revealing a selective role for presynaptic NMDARs in synaptic regulation. By dissociating presynaptic from postsynaptic receptor pools, our work explains earlier conflicting results and establishes presynaptic NMDARs as context-dependent but genuine contributors to cortical synaptic signaling.

### An NMDAR-mediated double dissociation in STDP

We uncovered a double dissociation between presynaptic and postsynaptic NMDAR function in STDP. tLTD required presynaptic but not postsynaptic NMDARs, whereas tLTP showed the reciprocal dependence, requiring postsynaptic NMDARs for induction while tolerating presynaptic deletion. This dissociation places NMDAR location at the center of STDP and refines classical STDP frameworks by separating the mechanisms underlying synaptic weakening and strengthening.^10,11,48,62^

For tLTD, our results align with recent work showing that presynaptic NMDARs drive L5 PC → PC depression.^47^ In that framework, presynaptic NMDARs decode post-before-pre spike timing via non-ionotropic signaling pathways, whereas the requirement for postsynaptic receptors remains uncertain. Earlier observations that tLTD persists with weak postsynaptic depolarization are consistent with this model.^63^ By selectively deleting NMDARs from defined synaptic compartments, we directly confirm these findings and resolve the locus of induction. Additionally, we show that postsynaptic NMDARs are not required for tLTD.

By contrast, tLTP depended on postsynaptic NMDARs for induction, in accordance with Hebbian coincidence-detection models.^1,60–62^ Although tLTP expression engaged both pre- and postsynaptic mechanisms, consistent with mixed loci of expression,^54,64^ presynaptic changes emerged only downstream of intact postsynaptic NMDARs.

Thus, while the canonical view of NMDARs as postsynaptic coincidence detectors^1,2^ captures part of STDP, our results argue against a unitary NMDAR-based induction model. Instead, our findings support a compartment-specific STDP framework in which temporal order is parsed by distinct NMDAR pools, with presynaptic and postsynaptic receptors participating in separable coincidence detectors for synaptic weakening and strengthening, respectively.^65^ Such a separation between induction and expression is compatible with theoretical work showing that mixed pre- and postsynaptic expression expands the computational repertoire of STDP beyond what either locus alone can support.^66,67^

### NMDARs and the link between synaptic and structural plasticity

Our results reveal a close association between NMDAR signaling and the long-term structural organization of cortical circuits. While synaptic plasticity may operate on timescales of minutes to hours,^62^ structural changes can emerge more slowly and span multiple scales, from dendritic spine remodeling over days to weeks^68^ to longer-term refinement of axonal and dendritic arbors.^58,59,69^ In this context, our findings suggest that NMDARs couple activity-dependent synaptic plasticity to the longer-term preservation of circuit architecture, consistent with evidence that NMDAR perturbations alter spine dynamics and neural morphology.^4^

In the global NMDAR deletion condition, basal dendritic spine density was reduced early, particularly on distal basal dendrites that host the majority of L5 PC → PC synapses.^56^ By contrast, sparse postsynaptic NMDAR deletion did not immediately affect overall spine density in juvenile animals, despite clear impairments in tLTP. Instead, spine-density deficits emerged gradually and became apparent only at later developmental stages. This temporal dissociation supports the idea that postsynaptic NMDAR signaling is dispensable for initial spine formation but required for the long-term stabilization or maintenance of excitatory synaptic contacts, consistent with evidence that activity-dependent plasticity primarily governs spine persistence rather than spine genesis,^4,68^ although compensatory changes in receptor composition cannot be excluded.

The delayed spine-density phenotype following sparse NMDAR deletion mirrors the functional dissociation observed in STDP: postsynaptic NMDARs were required for tLTP induction but dispensable for tLTD, whereas presynaptic NMDARs showed the opposite dependence. Notably, only loss of postsynaptic NMDARs — the receptors that govern associative synaptic strengthening — was accompanied by spine loss and altered dendritic architecture. This correspondence is consistent with a relationship between Hebbian potentiation and long-term synaptic stabilization.^4,6,7,68^

Our data do not support a direct instructive mapping between synaptic and structural plasticity. Rather, NMDAR signaling may define permissive conditions that favor long-term stabilization of strengthened synapses and loss of weakened connections. This view links synaptic and structural plasticity through NMDARs and shared activity dependence across timescales.

### NMDARs shape circuit architecture beyond synaptic strength

Beyond regulating synaptic efficacy, NMDAR signaling contributes to the architectural organization of cortical circuits. L5 PCs possess highly structured axonal and dendritic arbors that support recurrent intracortical and interlaminar connectivity.^56,57^ Consistent with this organization, we found that NMDAR deletion produced compartment- and layer-specific effects on neuronal morphology, preferentially altering axonal branching and proximal dendritic structure rather than uniformly affecting dendritic growth.^58^ Notably, these branching phenotypes did not track the changes we observed in synaptic release or STDP, consistent with longer-timescale roles of NMDARs in synapse stabilization and circuit maintenance.

Our data align with classical work linking neuronal activity and NMDAR signaling to dendritic growth and structural refinement.^58,59,69^ Consistent with emerging work, pre- and postsynaptic NMDAR perturbations can produce distinct effects on circuit architecture, supporting a compartment-specific organization of NMDAR function.^70^

By selectively reshaping axonal and dendritic domains engaged in recurrent excitation, NMDARs link synaptic plasticity to circuit-level organization in a compartment-specific manner. Mirroring the double dissociation observed in STDP, presynaptic and postsynaptic NMDARs exert distinct influences on neuronal structure, biasing where synaptic inputs are incorporated. Because dendritic geometry constrains integration, input selectivity, and synaptic plasticity,^71,72^ these effects position NMDARs as key regulators of cortical circuit architecture, rather than simple mediators of synaptic efficacy.

### Methodological implications for studying NMDAR function

This study highlights the importance of compartment-specific genetic deletion for dissociating presynaptic from postsynaptic NMDAR function. By enabling causal assignment of receptor signaling to defined synaptic compartments, this approach avoids confounds inherent to global manipulations.

Our findings also underscore that pharmacological approaches alone can yield ambiguous compartment-specific conclusions. Competitive antagonists act broadly, channel pore blockers incompletely suppress NMDAR signaling^39^ and spare non-ionotropic modes^31–38,47^, and calcium imaging preferentially reports ionotropic activity, yet some subunits and conditions limit Ca^2+^ influx.^40,73,74^

By combining ultrastructural localization with sparse genetic perturbation and direct physiological readouts, our approach provides a general framework for resolving receptor function in complex, recurrent circuits where signaling is spatially segregated or noncanonical. Together, these results underscore that resolving NMDAR function in cortical circuits requires approaches that distinguish pre- and postsynaptic compartments.

### Caveats and limitations

While our findings delineate distinct roles for pre- and postsynaptic NMDARs in synaptic and structural plasticity, our study focuses on compartmental receptor pools and does not distinguish synaptic from extrasynaptic NMDARs,^75,76^ or directly separate ionotropic from non-ionotropic signaling mechanisms.^25^ Moreover, deletion of the obligatory GluN1 subunit removes all NMDAR subtypes within a compartment, precluding subunit-specific conclusions.^30^

Unlike postsynaptic NMDARs, presynaptic NMDARs are not uniformly distributed across cortical synapses, but are restricted to specific synapse types.^43,77–79^ Consequently, the compartment-specific logic identified here may not generalize across all synapses. Presynaptic NMDARs are also developmentally regulated, but we focused on juvenile and adolescent cortex, where preNMDAR signaling has been implicated in critical-period plasticity.^79,80^

Because genetic deletion is chronic, eliminating NMDAR-mediated synaptic transmission may alter overall circuit drive,^81^ and could engage homeostatic adaptations in excitability and synaptic strength.^82^ However, our sparse deletion strategy argues against a purely global network explanation.

Future temporally controlled loss-of-function and rescue experiments will establish whether this compartment-specific logic extends to adult cortex and distinguish acute NMDAR signaling from longer-term compensation.

### Implications for cortical learning and disease-relevant plasticity

Our findings suggest that compartmentalized NMDAR signaling supports cortical learning by separating synaptic weakening and strengthening across presynaptic and postsynaptic receptor pools. This organization may allow circuits to remain adaptable while preserving recurrent structure.^62^

Such compartmentalization may also be relevant for disorders associated with NMDAR dysfunction, which involve both altered plasticity and circuit instability. Rather than uniform changes in NMDAR activity, imbalances between pre- and postsynaptic signaling could contribute to these phenotypes.^83^

More broadly, these results highlight a limitation of globally targeting NMDARs. Because distinct receptor pools support different forms of plasticity, non-selective modulation may disrupt circuit balance, underscoring the importance of considering NMDAR compartmentalization. Understanding how distinct NMDAR pools coordinate plasticity across timescales may therefore be essential for linking synaptic mechanisms to stable yet adaptable cortical computation.

## ACKNOWLEDGMENTS

We thank Alanna Watt, Keith Murai, Charles Bourque, Ed Ruthazer, Lina Dulgher and members of the Sjöström lab for helpful discussions. S.R. was supported by doctoral awards from the Fonds de recherche du Québec – Santé (FRQS; 317516), the Healthy Brains for Healthy Lives (HBHL) initiative, and the Research Institute of the McGill University Health Centre (RI-MUHC). K.L. was supported by an IBRO Summer Studentship. S.A. was supported by doctoral awards from the Fonds de recherche du Québec – Nature et technologies (FRQNT; 360969), HBHL, and the RI-MUHC. A.T. was supported by a Marie Skłodowska-Curie Fellowship (892837). R.L. was supported by a grant PID2024-155887OB-I00 funded by MCIN/AEI/ 10.13039/501100011033 and by “ERDF A way of making Europe”, by a grant from Castilla-La Mancha Regional Government and the European Regional Development Fund (SBPLY/24/180225/000007) and Universidad de Castilla-La Mancha (2025-GRIN-38362). P.J.S. was supported by the Canada Foundation for Innovation (LOF 28331), the Canadian Institutes of Health Research (PG 156223, 191969, 191997), the Fonds de recherche du Québec – Santé (CB 254033), the Natural Sciences and Engineering Research Council of Canada (DG/DAS 2017-04730, 2017-507818, 2024-06712), and the Donald S. Wells Distinguished Scientist Award.

## AUTHOR CONTRIBUTIONS

S.R. and P.J.S. conceived and designed the study. S.R. performed electrophysiological and imaging experiments, with A.T. contributing to a subset of recordings. K.L., C.B.K., N.C., V.Y.L., and G.M. performed morphometric analyses, including cell and spine quantification. S.A. performed linear mixed-effects modeling and statistical analyses. R.L. conducted immuno-electron microscopy experiments and particle quantification. P.J.S. supervised the project and developed custom software for data acquisition and analysis. S.R. and P.J.S. analyzed the data and wrote the manuscript.

## STAR⍰METHODS

- KEY RESOURCES TABLE
- RESOURCE AVAILABILITY

`o` Lead contact
`o` Materials availability
`o` Data and code availability
- EXPERIMENTAL MODEL AND SUBJECTS DETAILS

`o` Mouse strains and husbandry
`o` Neonatal injections
- METHOD DETAILS

`o` Electron Microscopy
`o` Quantitative Immunogold Analysis
`o` Slice Preparation and Basic Electrophysiology
`o` Pharmacology
`o` Evoked Release and Long-Term Plasticity
`o` Spontaneous Release
`o` 2-Photon Laser Scanning Microscopy and Uncaging
`o` Histology and Morphometry
- QUANTIFICATION AND STATISTICAL ANALYSIS

## STAR⍰Methods

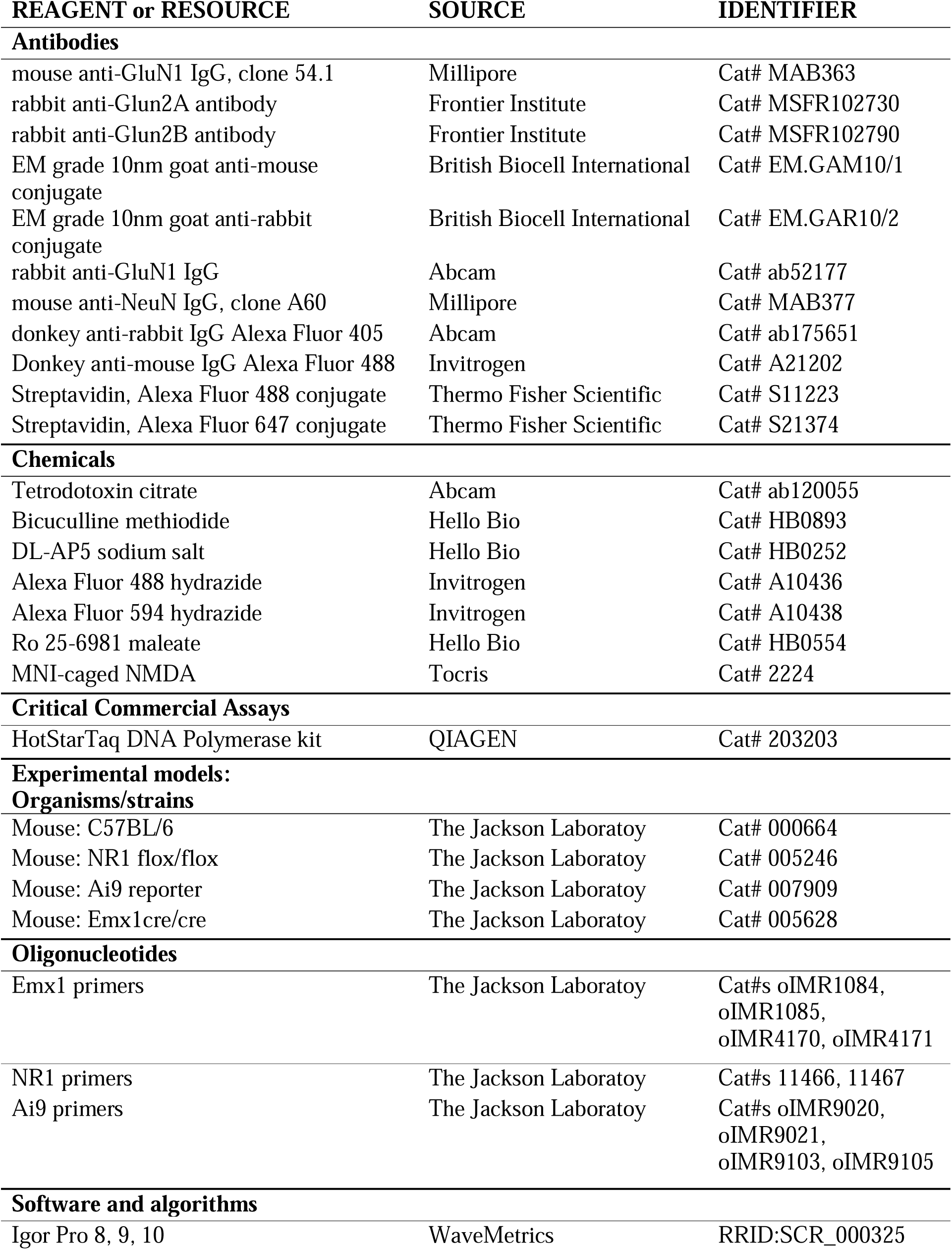
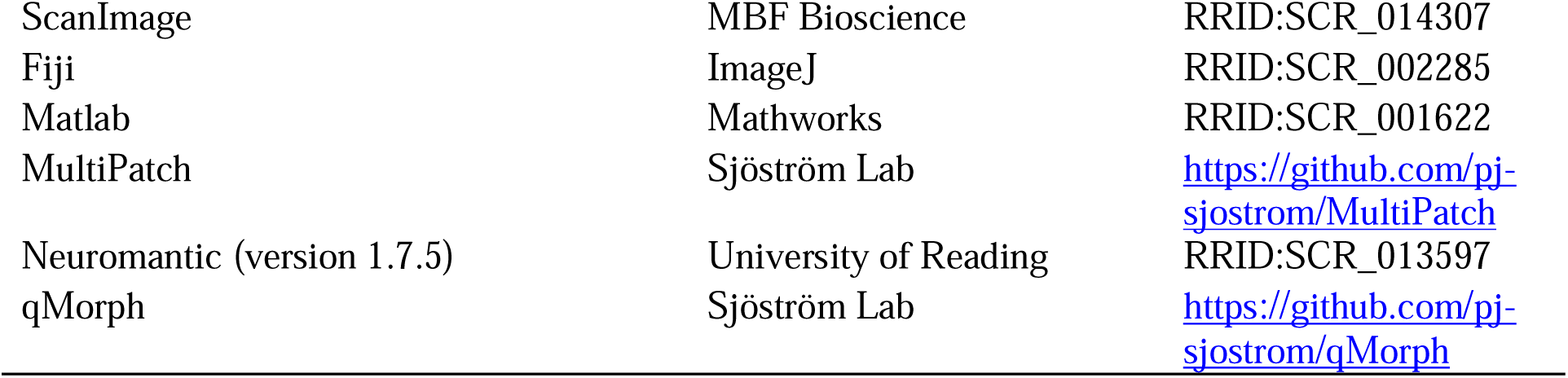
KEY RESOURCES TABLE.

## RESOURCE AVAILABILITY

### Lead contact

Further information and requests for resources and reagents should be directed to and will be fulfilled by the lead contact, Jesper Sjöström.

### Materials availability

This study did not generate new unique reagents.

### Data and code availability

- All data reported in this paper will be available on Dryad at https://doi.org/10.5061/dryad.ht76hdrx6
- All custom code used for data acquisition and data analysis are available from the lead contacts on reasonable request, as well as in GitHub links provided in the Key Resource Table section.
- Any additional information required to reanalyze the data reported in this paper is available from the lead contact upon request.

## EXPERIMENTAL MODEL AND SUBJECT DETAILS

### Mouse strains and husbandry

Animal procedures conformed to the Canadian Council on Animal Care as overseen by the Montreal General Hospital Facility Animal Care Committee, with appropriate licenses. C57BL/6J (Strain 000664), homozygous Emx1cre/cre (005628), homozygous NR1^flox/flox^ (005246), and Ai9 reporter animals (007909) were obtained from The Jackson Laboratory. Global NMDAR deletion Emx1^cre/+^;NR1^flox/flox^;Ai9^tdTomato/+^ animals and Emx1^cre/+^;NR1^flox/flox^;Ai9^tdTomato/+^ (“WT”) littermates were generated by crossing Emx1^cre/cre^ mice with NR1^flox/flox^ and Ai9 reporter animals. Genotyping was carried out using standard methodology with Jackson Laboratory primers (Emx1: oIMR1084, oIMR1085, oIMR4170, oIMR4171; NR1: 11466, 11467; Rosa26: oIMR9103, oIMR9105, oIMR9020, oIMR9021) using QIAGEN HotStarTaq DNA Polymerase kit (203203) and dNTPs from Invitrogen/Thermo-Fisher (18427-013). All transgenic animals apart from Emx1^cre/+^;NR1^flox/flox^;Ai9^tdTomato/+^ mice had no abnormal phenotype. Emx1^cre/+^;NR1^flox/flox^;Ai9^tdTomato/+^ mice displayed developmental disruption, and thus, we set the end point for these animals to weaning age (P20-22).

The C57BL/6 mice used in the immuno-EM experiments (see below) were obtained from the Animal House Facility of the School of Medicine of the University of Castilla-La Mancha. Care and handling followed Spanish and European Union regulations, as approved by the institutional Animal Care and Use Committee.

All mice were group housed in a standard 12 hour light/12 hour dark cycle with food and water available *ad libitum*.

### Neonatal injections

NR1^flox/flox^ pups P0-2 were injected with AAV-eSYN-mCherry-T2A-iCre-WPRE (VB4856, Vector Biolabs) or AAV-eSYN-EGFP-T2A-iCre-WPRE (VB4855, Vector Biolabs) into V1 using the following coordinates (in mm, relative to lambda): medial-lateral ± 1.10; anterior-posterior 0. Pups were placed on ice for hypothermic anesthesia before fixing the head with ear bars to perform three infusions 0.2-0.3µL of a 1:10 viral dilution in phosphate-buffered saline (PBS) at depths of 0.20 mm, 0.15 mm, and 0.10 mm below the pial surface, at a rate of 0.25 µL/minute with a 33-gauge needle attached to a 10 µL gas-tight syringe (Hamilton Instruments, Reno, NV).

## METHOD DETAILS

### Electron Microscopy

Immuno-EM experiments were performed in the laboratory of R.L. using established post-embedding immunogold protocols.

A monoclonal antibody against GluN1 (clone 54.1 MAB363) was obtained from Millipore (Germany). This antibody was directed against a fusion protein corresponding to amino acid residues 660–811, representing the extracellular loop between transmembrane regions III and IV of the NR1 subunit.^84^ The specificity of this antibody has been previously validated.^84^ An affinity-purified polyclonal rabbit anti-GluN2A (MSFR102730) and an affinity-purified polyclonal rabbit anti-GluN2B (MSFR102790) were obtained from Frontier Institute (Hokkaido, Japan).

The secondary antibodies used were as follows: goat anti-mouse IgG conjugated to 10 nm gold particles and goat anti-rabbit IgG conjugated to 10 nm gold particles (1:100; British Biocell International, Cardiff, UK).

Immunohistochemical reactions at the electron microscopic level were carried out using the post-embedding immunogold method as described earlier.^85^ Briefly, animals (n = 3 male P21 C57BL/6 mice) were anesthetized by intraperitoneal injection of ketamine-xylazine 1:1 (0.1 mL/kg b.w.) and transcardially perfused with ice-cold fixative containing 4% paraformaldehyde, 0.1% glutaraldehyde and 15% saturated picric acid solution in 0.1 M phosphate buffer (PB) for 15 min. Vibratome sections of 500 μm thickness were placed into 1 M sucrose solution in 0.1 M PB for 2 h before they were slammed on a Leica EM CPC apparatus. Samples were dehydrated in methanol at -80°C and embedded by freeze-substitution (Leica EM AFS2) in Lowicryl HM 20 (Electron Microscopy Sciences, Hatfield, USA), followed by polymerization with UV light. Then, ultrathin 80-nm-thick sections from Lowicryl-embedded blocks of the visual cortex were picked up on coated nickel grids and incubated on drops of a blocking solution consisting of 2% human serum albumin (HSA) in 0.05 M TBS and 0.03% Triton X-100. The grids were incubated with GluN1, GluN2A or GluN2B antibodies (10 μg/mL in 0.05 M TBS and 0.03% Triton X-100 with 2% HSA) at 28°C overnight. The grids were incubated on drops of goat anti-rabbit IgG conjugated to 10 nm colloidal gold particles (1:100; British Biocell International, Cardiff, UK) in 2% HSA and 0.5% polyethylene glycol in 0.05 M TBS and 0.03% Triton X-100. The grids were then washed in TBS and counterstained for electron microscopy with 1% aqueous uranyl acetate followed by Reynolds’s lead citrate. Ultrastructural analyses were performed in a JEOL JEM-1400 Flash electron microscope.

### Quantitative Immunogold Analysis

The quantifications were performed in three animals. Morphologically identifiable excitatory synapses containing dendritic spines with a clear PSD and axon terminals with presynaptic active zone and/or clear vesicles were assessed. Only morphologically identifiable excitatory synapses that contained at least one immunogold particle were included for quantification. All immunogold particles associated with included synapses were quantified. Then, the axial distribution of GluN1, GluN2A and GluN2B across layer 5 excitatory synapses was determined. The distance from the border between the plasma membrane of the PSD (0 nm) to the center of each gold particle was recorded along an axis perpendicular to the postsynaptic membrane. Positive values indicate postsynaptic direction, and negative values indicate presynaptic direction. Relative distance of the immunoparticles to postsynaptic membrane structure was allocated to 30 nm wide bins and expressed as percentage.

### Slice Preparation and Basic Electrophysiology

All animal procedures conformed to the Canadian Council on Animal Care as overseen by the Montreal General Hospital Facility Animal Care Committee, with appropriate licenses. Male or female P10-20 mice were anaesthetized with isoflurane and sacrificed once the limb withdrawal reflex was determined absent.

Mouse brains were dissected in ice-cold (<4L) artificial cerebrospinal fluid (ACSF: 125 mM NaCl, 2.5 mM KCl, 1.25 mM NaH_2_PO_4_, 26 mM NaHCO_3_, 2 mM CaCl_2_, 1 mM MgCl_2_, 25 mM D-Glucose, bubbled with 95% O_2_/5% CO_2_, osmolality adjusted to 338 mOsm/kg with D-glucose). Using a Campden Instruments 5000mz-2 vibratome, brain slices of 300 µm were cut from the visual cortex at angle normal to the pial surface to maximally preserve apical dendrites and principal axons. Slices were transferred to incubation chamber at 32L for 20 min before cooling down to room temperature (∼23L). Experiments were performed with ACSF perfused at 32-34L, maintained by a resistive inline heater (Scientifica, or Warner Perfusion heater 640102 with controller 642400).

Patch pipettes of 4-6 MΩ resistance were pulled from standard-wall borosilicate capillaries (1.5 mm outer diameter, 0.86 mm inner diameter, either Harvard Apparatus G150F-4, or Sutter Instrument BF150-86-10) with a P-1000 electrode puller (Sutter Instruments). Pipettes were loaded with internal solution (5 mM KCl, 115 mM K-Gluconate, 10 mM HEPES pH 7.4 calibrated with KOH, 4 mM Mg-ATP, 0.3 mM Na-GTP, 10 mM Na-phosphocreatine, 0.1% w/v Biocytin, adjusted with KOH to pH 7.4, osmolality adjusted to 310 mOsm/kg with sucrose). For 2-photon laser-scanning microscopy, internal solution was supplemented with 20-150 µM Alexa Fluor 488 or Alexa Fluor 594 (Life Technologies).

Whole-cell recordings were performed using BVC 700A (Dagan) or MultiClamp 700B amplifiers (Molecular Devices). Current clamp recordings were filtered at 20 kHz and acquired at 40 kHz using PCI-6229 or PCIe-6323 boards (National Instruments) with custom-written software (https://github.com/pj-sjostrom/MultiPatch) running in Igor Pro 7, 8 or 9 (WaveMetrics). Series resistance, input resistance, resting membrane potential or holding current, excitatory postsynaptic potential (EPSP), and perfusion temperature were monitored online and assessed offline. Series resistance was continuously monitored but not compensated. Recordings were not adjusted for liquid junction potential (∼10 mV).

Neurons were patched with an Olympus LUMPlanFL N 40×/0.80 objective using infra-red video Dodt contrast. Primary visual cortex was identified based on the presence of layer 4. L5 PCs were targeted for recording based on their characteristically large pyramid-shaped somata and thick apical dendrites.

### Pharmacology

To isolate AMPAR-mediated miniature excitatory postsynaptic currents (mEPSCs), tetrodotoxin citrate (TTX; 0.1 µM; Abcam, #ab120055) and bicuculline methiodide (30 µM; Hello Bio, #HB0893) were added to the bath. For wash-in experiments, NMDARs were blocked using DL-AP5 sodium salt (200 µM; referred to as AP5; Hello Bio, #HB0252) or the GluN2B-selective antagonist Ro 25-6981 maleate (0.5 µM; referred to as Ro; Hello Bio, #HB0554). Uncaging was performed by locally puffing 1 mM 4-Methoxy-7-nitroindolinyl-caged N-methyl-D-aspartic acid (MNI-NMDA; Tocris #2224), diluted in ACSF.

### Evoked Release and Long-Term Plasticity

Because connectivity between L5 PCs is sparse, we performed quadruple whole-cell recordings, allowing simultaneous testing of up to 12 possible connections per quadruple attempt.^86^ Seals were formed on four neurons and ruptured in rapid succession. Cells were targeted based on the fluorescent tag of the virus (mCherry or EGFP) as visualized by two-photon microscopy. In each quadruple recording, at least one PC expressed the fluorescent tag and at least one did not. To ensure reliable detection of fluorescent tag, we avoided patching cells located deep into the acute slice.

To test for connections, five spikes were evoked at 30 Hz by 5-ms long current injections (∼1.3 nA) every 20 s for at least ten repetitions. Spikes in different cells were temporally offset by 700 ms to avoid inadvertent plasticity induction. If no EPSPs were detected, recordings were terminated and new cells were patched.

For evoked release experiments with NMDAR blockade, five-spike bursts at 30 Hz were delivered every 20 s. After a stable baseline (10-15 min), AP5 was washed in for 10 min while stimulation continued for up to 200 repetitions, or 67 min in total, similar to our prior studies.^37,38,42,43^ Slices were replaced after each AP5 experiment.

For tLTD experiments, baseline activity was reduced to avoid accidental rundown in Del → WT or Del → Del pairs akin to AP5 wash-in above, so consisted of single spikes every 10 s, with induction consisting of five post-before-pre spikes (Δt = -25 ms) delivered at either 1 Hz or 20 Hz every 10 s for 15 repetitions.^10,42,47^ The current tLTD dataset expands on a previously published dataset showing a need for presynaptic NMDARs.^47^ For tLTP experiments, baseline was likewise reduced to paired spikes every 15 s, with induction consisting of five pre-before-post spikes (Δt = +10 ms) delivered at 50 Hz every 10 s for 15 repetitions.^10,47^ Slices were fixed immediately after recordings, see Histological Preparation below.

Across all experiments, input resistance, resting membrane potential, EPSP amplitude, and bath temperature were continuously monitored. Recordings were discarded or truncated if input resistance changed by >30%, membrane potential shifted by >8 mV, or bath temperature deviated from 32–34 °C. Baseline periods were considered unstable if EPSP amplitude showed a significant monotonic trend over time, assessed by a Pearson correlation test (p < 0.05), and such recordings were excluded from further analysis.

### Spontaneous Release

Spontaneous excitatory synaptic transmission was recorded in voltage clamp in the presence of tetrodotoxin (TTX) and bicuculline, as described above. For sparse NMDAR deletion experiments, recordings were obtained at a holding potential of −80 mV. For global NMDAR deletion experiments, recordings were performed at −70 mV using ACSF containing reduced MgCl_2_ (0.2 mM).

Miniature EPSCs (mEPSCs) were acquired in 25-s sweeps every 30 s and low-pass filtered offline at 2 kHz. Individual events were automatically detected using in-house software^87^ implemented in Igor Pro 9, with detection criteria including amplitude >5 pA and rise time <3 ms. Overlapping events and unstable baselines were automatically excluded, and sweeps containing large artifacts (e.g., electrical noise) were manually discarded.

As for evoked release experiments, input resistance and series resistance were monitored throughout using a −5 mV, 250-ms-long test pulse. Recordings were rejected if series resistance exceeded 40 MΩ or changed by >20%. Additional exclusion criteria matched those used for evoked release: recordings were discarded if input resistance or membrane potential stability criteria were violated, or if baseline mEPSC frequency exhibited a significant monotonic trend over time (Pearson correlation test, p < 0.05). Recordings shorter than 20 min following drug wash-in were excluded. For wash-out experiments, mEPSC frequency was additionally required to recover to within 30% of baseline.

For baseline comparisons of spontaneous release (Table S1), all recordings that met electrophysiological quality criteria were included, regardless of initial mEPSC frequency. For drug wash-in/wash-out time-course analyses, recordings with an initial mEPSC frequency <1 Hz were excluded to ensure reliable estimation of frequency changes. Ensemble time courses (e.g., Figures 2D and 2I) were normalized to the pre-drug baseline period.

### Two-Photon Laser Scanning Microscopy and Uncaging

Two-photon laser scanning microscopy was performed using imaging workstations custom-built around either a BX51WI microscope (Olympus) or a SliceScope microscope (Scientifica Ltd., UK), as previously described.^43^ Fluorescence detection was achieved using R3896 bialkali photomultiplier tubes (PMTs; Hamamatsu), mounted either in epifluorescence or substage configurations. Laser scanning was performed with 3-mm 6215H galvanometric mirrors (Cambridge Technology).

For uncaging experiments, fluorescence was collected through the condenser to allow delivery of the 405-nm laser beam to the acute slice; otherwise, fluorescence was collected through the objective. Two-photon excitation was provided by a MaiTai HP titanium–sapphire laser (Spectra-Physics) tuned to 750 nm, 820 nm, or 920 nm. Lasers were gated using either an SH05/SC10 shutter–controller pair (Thorlabs) or Uniblitz LS6ZM2 shutters driven by a VCM-D1 controller (Vincent Associates, Rochester, NY, USA). Laser power was manually attenuated using a polarizing beam splitter in combination with a half-wave plate (Thorlabs GL10-B and AHWP05M-980). Laser output power was monitored using a PM100A power meter equipped with an S121C sensor (Thorlabs).

Fluorescence was collected through an Olympus 40× objective (NA0.8) or condenser (NA1.4) and separated using either an FF665-Di01 or FF695-Di01 dichroic mirror (Semrock). Green and red fluorescence signals were further separated using an FF560-Di01 dichroic beam splitter (Semrock), together with an ET525/50m green emission filter and an ET630/75m red emission filter (Chroma). Laser-scanning Dodt gradient contrast images were obtained by collecting transmitted laser light after the spatial filter with an amplified photodiode (Thorlabs PDA100A-EC).

Imaging data were acquired using ScanImage software,^88^ versions 3.7 (customized), 2019, 2020, or 2021, running in MATLAB (MathWorks), interfaced via PCI-6110 or PCIe-6374 data acquisition boards (National Instruments).

Following whole-cell recordings, L5 pyramidal cell morphologies were acquired as stacks of 512×512-pixel images with z-steps of 2 µm. Each optical section was averaged from three frames acquired at 2-4 ms per line. Typically, four image stacks were required to capture the complete dendritic and axonal arbor of each neuron. For NMDAR deletion experiments, stacks used to assess mCherry, EGFP or tdTomato fluorescence were acquired prior to approaching the slice with recording electrodes.

The uncaging laser was implemented as previously described.^89^ Briefly, a 405-nm solid-state laser diode (Amazon.ca, output ∼120 mW) was gated using a Uniblitz LS3ZM2 shutter driven by a VCM-D1 controller (Vincent Associates) with a pulse duration of 10 ms. The uncaging beam was combined with the Ti:Sa two-photon imaging beam using an FF665-Di02 dichroic mirror (Semrock). Both imaging and uncaging beams were steered using the same pair of 3-mm 6215H galvanometric mirrors (Cambridge Technology). The power of the 405-nm laser beam measured at the objective back aperture was approximately 10 mW.

### Histology and Morphometry

Following electrophysiological recordings, acute brain slices were fixed overnight in 4% paraformaldehyde (PFA) and subsequently transferred to PBS. Neurons filled intracellularly with biocytin during recordings were visualized using streptavidin conjugated to Alexa Fluor 488 or Alexa Fluor 647 (Thermo Fisher Scientific), enabling three-dimensional reconstructions of axodendritic arbors as well as quantification of basal dendritic spine density.

For immunolabeling experiments, animals were anesthetized with isoflurane, exsanguinated by intracardial perfusion with PBS, and fixed with 4% PFA. Brains were removed and post-fixed in 4% PFA for 24 h, cryoprotected in PBS containing 30% sucrose for 48 h, embedded in optimal cutting temperature (OCT) compound (Thermo Fisher Scientific), and stored at -80 °C until sectioning. Coronal cryostat sections (45-50 µm) were immunostained for GluN1 (Abcam ab52177; 1:50 or 1:100) and NeuN (MilliporeSigma MAB377; 1:500), followed by incubation with appropriate secondary antibodies (Abcam ab175651, 1:1000; Invitrogen A21202, 1:1000). Sections were mounted on Superfrost Plus slides (Thermo Fisher Scientific) using ProLong Glass Antifade Mountant (Thermo Fisher Scientific).

Confocal imaging was performed on a Zeiss LSM780 laser-scanning confocal microscope using either a 20×/0.40 NA long-working-distance Plan-Neofluar objective or a 40×/1.20 NA water-immersion C-Apochromat objective with correction collar, controlled by ZEN 2010 software (Zeiss). Fluorophores were excited using 405-, 488-, 561-, or 633-nm laser lines as appropriate. Image brightness and contrast were adjusted in Fiji/ImageJ.^90^

Three-dimensional neuronal reconstructions were generated using Neuromantic software (version 1.7.5). Quantitative morphometric analyses were performed using the in-house qMorph software package (https://github.com/pj-sjostrom/qMorph) running in Igor Pro 9, as previously described.^91^ Reconstructions were derived either from 2-photon laser scanning microscopy images or from confocal stacks of biocytin-labeled neurons. As no left–right asymmetry was detected and to increase sampling, density maps (Figures 8A, S10A, S11A) were computed with each reconstruction in its original and mirrored orientations.

Basal dendritic spine density was quantified from confocal images of biocytin-stained slices acquired with a Zeiss Plan-Apochromat 63×/1.40 NA oil-immersion objective.

## QUANTIFICATION AND STATISTICAL ANALYSIS

All statistical analyses were performed in Igor Pro 9 (WaveMetrics) unless otherwise stated. Data are reported as mean ± SEM, and *n* denotes the number of cells or synaptic connections unless otherwise specified. Data from male and female animals were pooled.

Two-group comparisons were performed using Student’s *t* test. When variance heterogeneity was detected by an *F* test (*p* < 0.05), unequal-variance *t* tests were applied. For comparisons involving more than two groups, we used one-way ANOVA, or Welch’s ANOVA when Bartlett’s test indicated heteroscedasticity (*p* < 0.05), followed by Bonferroni-corrected post hoc pairwise comparisons when appropriate. In cases where normality was uncertain (e.g., tLTP in Figure 6), nonparametric Kruskal–Wallis and Mann–Whitney tests were additionally performed and yielded the same statistical conclusions.

Correlation analyses were performed using Pearson’s *r*, with significance assess by Student’s *t* test. In spontaneous release experiments, cumulative distributions were compared using the Kolmogorov–Smirnov (KS) test. For coefficient-of-variation (CV) analyses, Wilcoxon’s rank test was used to compare the angle φ relative to the unity diagonal (45°).

For Figure 1B, a chi-squared (χ^2^) test assessed whether GluN puncta were distributed differently between presynaptic and postsynaptic compartments, with expected values based on an equal (50/50) distribution. For Figure 1C, a Monte Carlo simulation (10,000 iterations) randomly reassigned GluN2A and GluN2B identities across the pooled set of 351 measured gold-particle locations (counts preserved: 174 GluN2A and 177 GluN2B) to estimate the probability of observing a postsynaptic GluN2A/GluN2B ratio at least as large as the measured value by chance.

Sholl analyses were evaluated using linear mixed-effects models (LMMs)^92^ implemented in RStudio (Posit Software), based on code originally developed by Adrian Gabriel Zucco (https://github.com/adrigabzu/sholl_analysis_in_R) using the *lme4* and *lmerTest* packages. Intersections were modeled as a function of radial distance, experimental condition, and their interaction, with random intercepts for individual neurons nested within slices and animals. Mixed-effects modeling was used because Sholl measurements at different radial distances from the same neuron are not independent. Fixed-effect coefficients were evaluated using *t* tests with Satterthwaite-approximated degrees of freedom. Planned directional tests of genotype effects across radii were corrected for multiple comparisons using the Benjamini–Hochberg false discovery rate procedure.

For Supplementary Figure S8, connectivity rates were compared using χ^2^ tests on 2×2 contingency tables testing hypothesized presynaptic versus postsynaptic effects on connectivity. Multiple comparisons were controlled using Bonferroni’s correction.

Box plots display the median and interquartile range (25^th^ - 75^th^ percentiles), with whiskers indicating the full data range, minimum to maximum. Significance levels are denoted as * for *p* < 0.05, ** for *p* < 0.01 and *** for *p* <0.001.

**Supplementary Figure S1:**
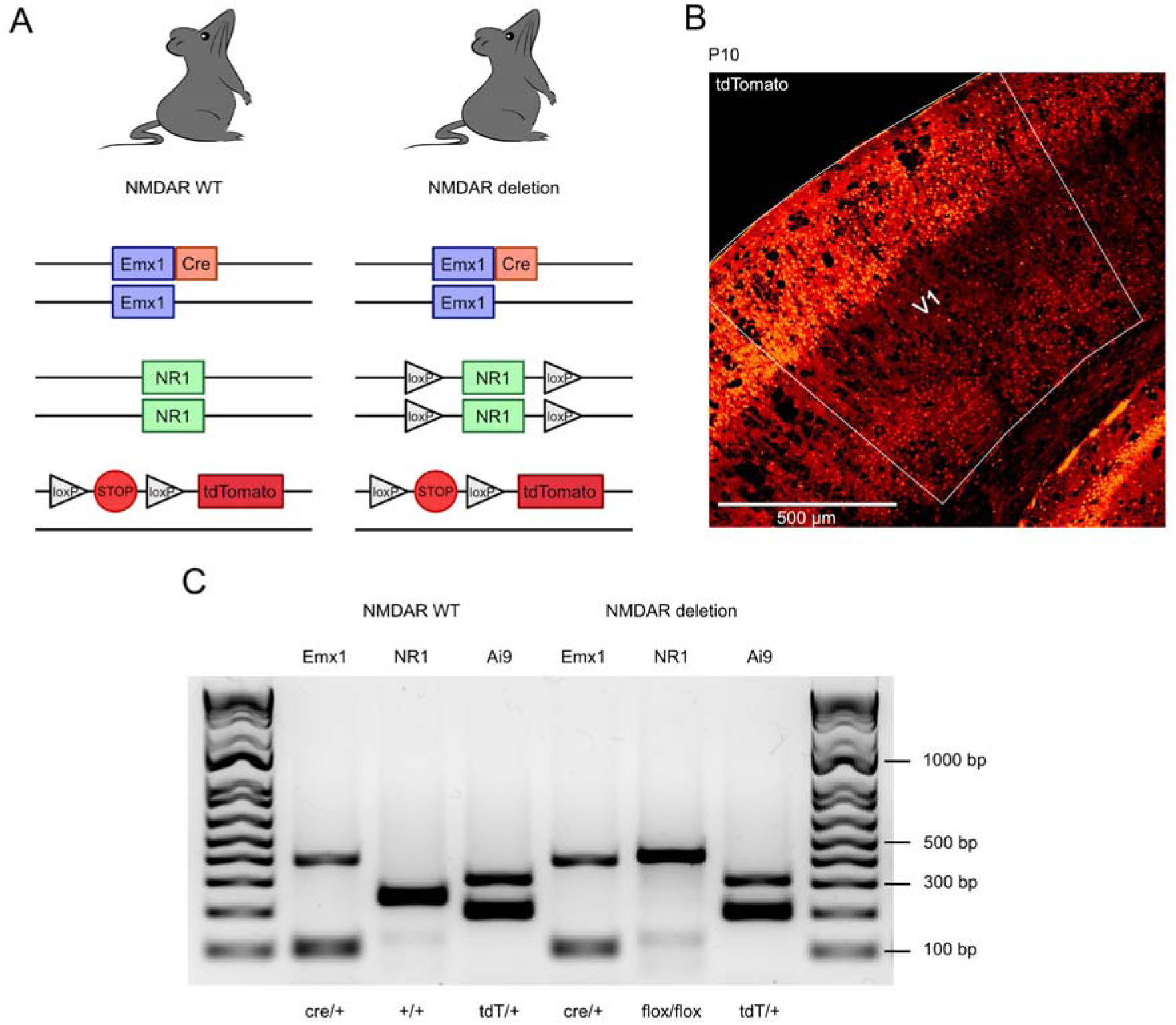
Emx1-Cre drives widespread cortical recombination in global NMDAR deletion mice, Related to Figure 2. (A) Schematic of Emx1, NR1, and Ai9 alleles in WT (Emx1^cre/+;^NR^+/+^;Ai9^tdTomato/+^) and global NMDAR deletion mice (Emx1^cre/+;^NR1^flox/flox^;Ai9^tdTomato/+^). (B) Representative confocal image of a coronal slice from a juvenile (P10) NMDAR deletion mouse showing widespread tdTomato fluorescence, indicating broad Emx1-Cre–mediated recombination across cortical layers. (C) Representative agarose gel confirming the presence of Emx1-Cre, NR1 floxed or WT alleles, and Ai9 tdTomato reporter in control and NMDAR deletion animals.

**Supplementary Figure S2:**
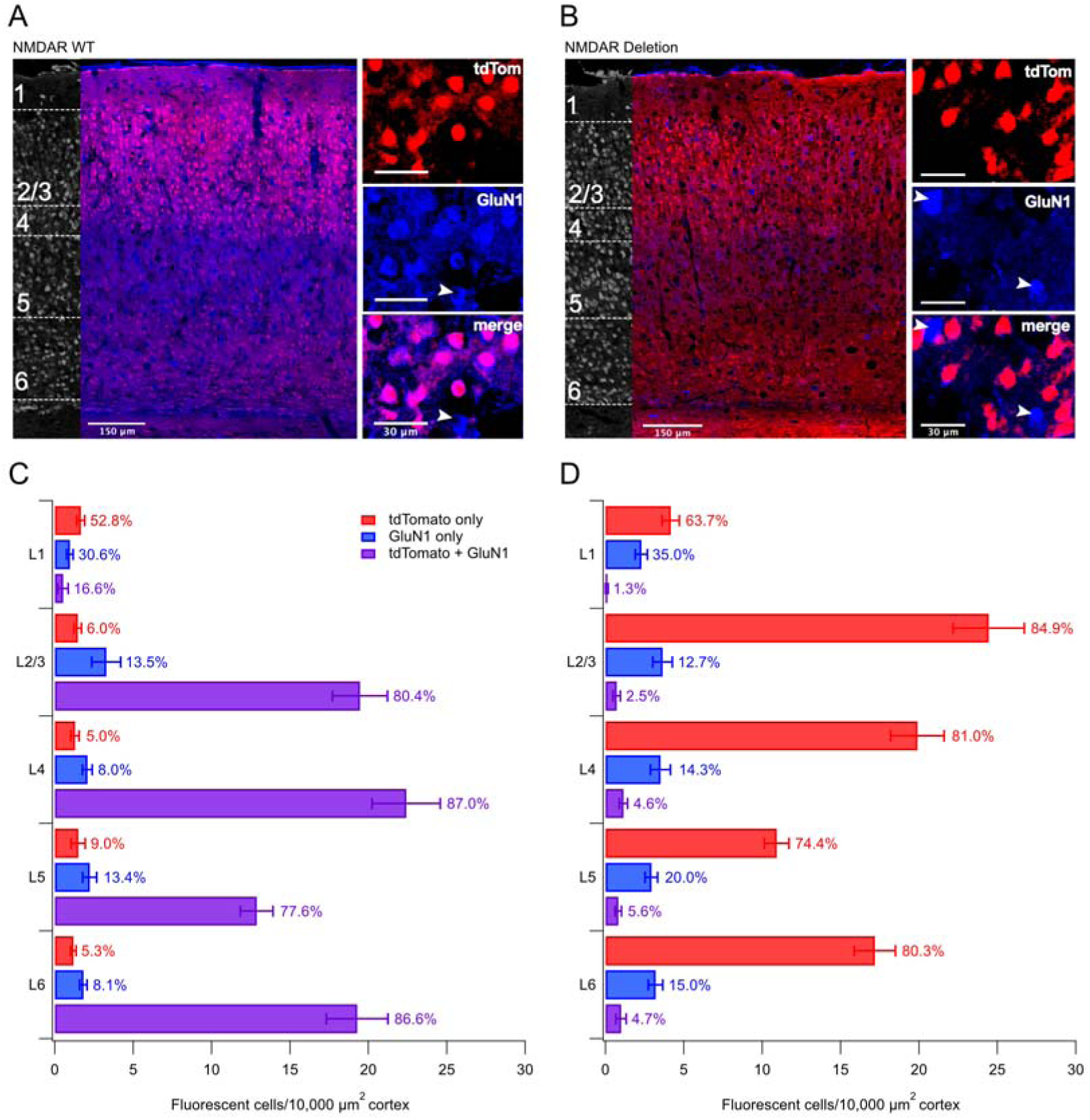
Global deletion removed NMDARs in tagged cells, Related to Figure 2. (A) Representative confocal image of a 50-μm-thick coronal brain section from an Emx1^cre/+^;NR1^+/+^;Ai9^tdTomato/+^ mouse (NMDAR WT), labelled for NeuN (grayscale, left), GluN1 (blue), and tdTomato (red). Arrowheads indicate example cells. (B) Same as in (A), but from an Emx1^cre/+^;NR1^flox/flox^;Ai9^tdTomato/+^ mouse (NMDAR Deletion). (C and D) NMDARs were efficiently deleted in tagged cells, as evidenced by colocalization in L5 being reduced from 89 ± 3% in NMDAR WT mice to 6 ± 1% in NMDAR Deletion animals (Mann-Whitney p < 0.001), where colocalization was calculated as the fraction of tdTomato-positive cells that were GluN1-positive. Although we focus on L5 here, NMDAR deletion was similarly efficacious in other cortical layers (see Dryad data set). Quantification of cell densities (cells per 10,000 μm^2^ cortex) expressing GluN1 only (blue), tdTomato only (red), or both GluN1 and tdTomato (purple) across cortical layers in NMDAR WT (*N* = 7 animals, 1–4 cortical sections per animal) and NMDAR Deletion mice (*N* = 4 animals, 3 cortical sections per animal). Percentage next to SEM bars denotes proportion of cells within the respective layer expressing either GluN1 only, tdTomato only, or both GluN1 and tdTomato.

**Supplementary Figure S3:**
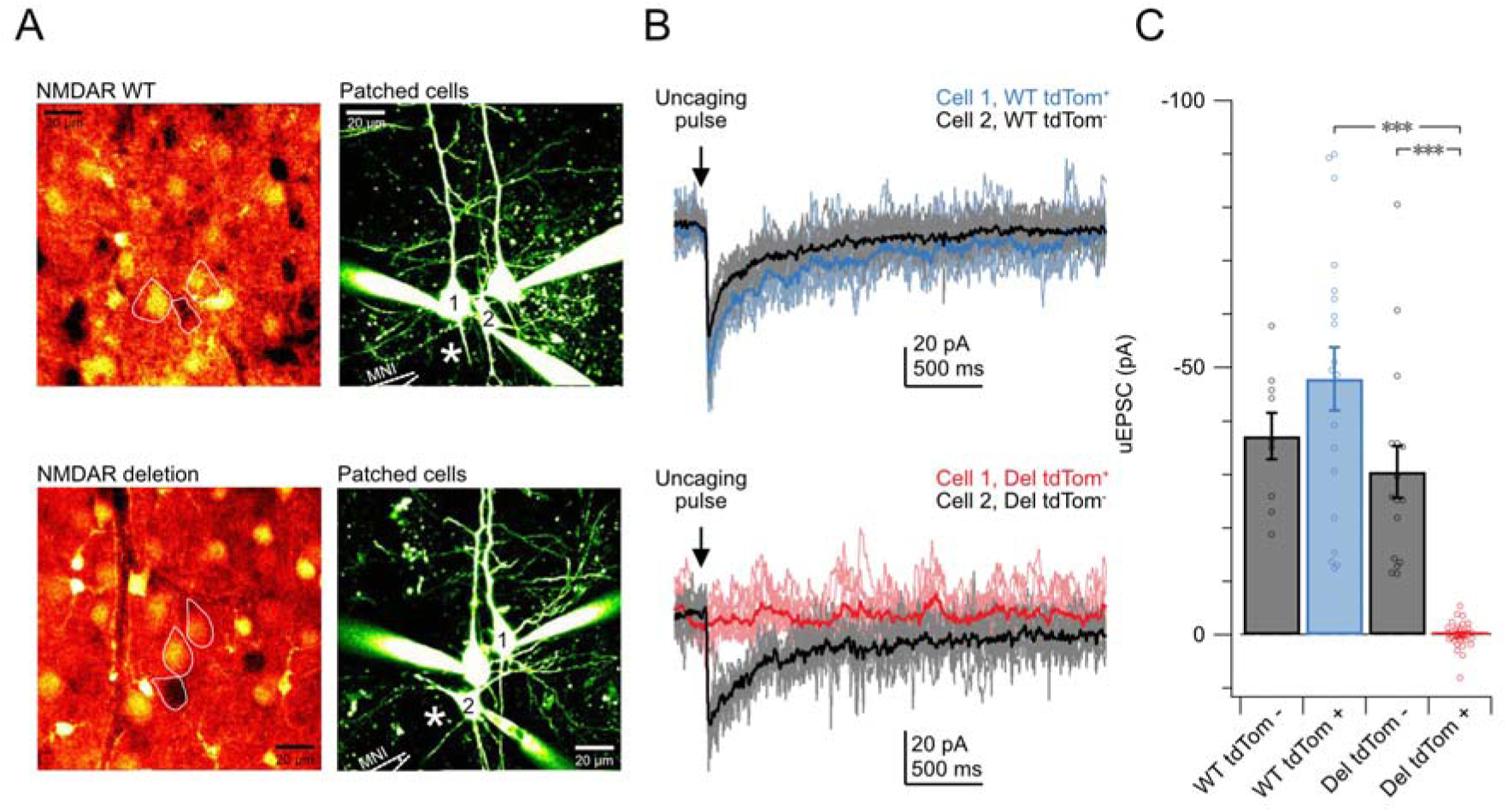
NMDA uncaging confirmed loss of functional NMDARs after global GluN1 deletion, Related to Figure 2. (A) Two-photon images of patched neurons in acute slices from a WT littermate (top, Emx1^cre/+;^NR^+/+^;Ai9^tdTomato/+^) and from a global NMDAR deletion mouse (bottom, Emx1^cre/+;^NR1^flox/flox^;Ai9^tdTomato/+^). Left: tdTomato fluorescence tagged recombined cells. Right: Sample 2-photon images of patched cells, with MNI-NMDA puff pipette and laser uncaging spot indicated (*). (B) Example uncaging-evoked EPSC traces evoked by laser uncaging of MNI-NMDA at the sites indicated in (A) (average of 10 sweeps). (C) Quantification of uncaging-evoked EPSCs confirmed efficient elimination of functional NMDARs in Cre-expressing pyramidal cells in the global deletion model. In NMDAR deletion animals, tdTomato-positive PCs (Del tdTomL, *n* = 31) showed no detectable responses to NMDA uncaging, in contrast to neighboring tdTomato-negative presumed interneurons (Del tdTomL, *n* = 16). In WT littermates, tdTomato-positive PCs (WT tdTomL, *n* = 19) responded robustly, with amplitudes indistinguishable from tdTomato-negative interneurons (WT tdTomL, *n* = 9). Student’s t test.

**Supplementary Figure S4:**
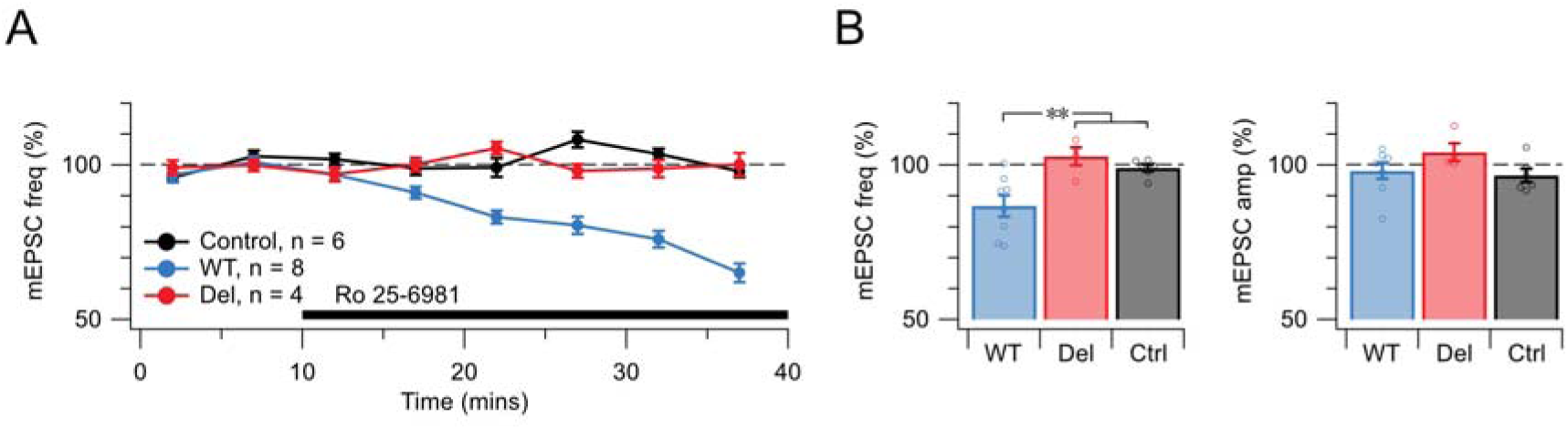
Spontaneous release regulation is most consistent with presynaptic GluN2B-containing NMDARs, Related to Figure 2. The GluN2B-selective NMDAR antagonist Ro 25-6981 significantly reduced mEPSC frequency without affecting mEPSC amplitude in WT animals (Welch’s ANOVA, p < 0.05; Bonferroni-corrected post hoc comparisons), indicating that GluN2B-containing NMDARs regulate spontaneous release. This effect mirrors prior findings demonstrating presynaptic NMDAR control of spontaneous transmission at V1 L5 PC → PC synapses.^37,38,42^ This effect of Ro 25-6981 is consistent with prior literature showing that pre- but not postsynaptic NMDARs at V1 L5 PC → PC synapses are sensitive to GluN2B-specific blockers at this age.^38,42,44^ However, in slices from Emx1^cre/+^; NR1^flox/flox^;Ai9^tdTomato/+^ mice (see Methods) with GluN1 globally deleted (“Del”), Ro 25-6981 had no effect and was indistinguishable from controls. Together, these results support the conclusion that spontaneous excitatory release onto V1 L5 pyramidal neurons is regulated by presynaptic, GluN2B-containing NMDARs.

**Supplementary Figure S5:**
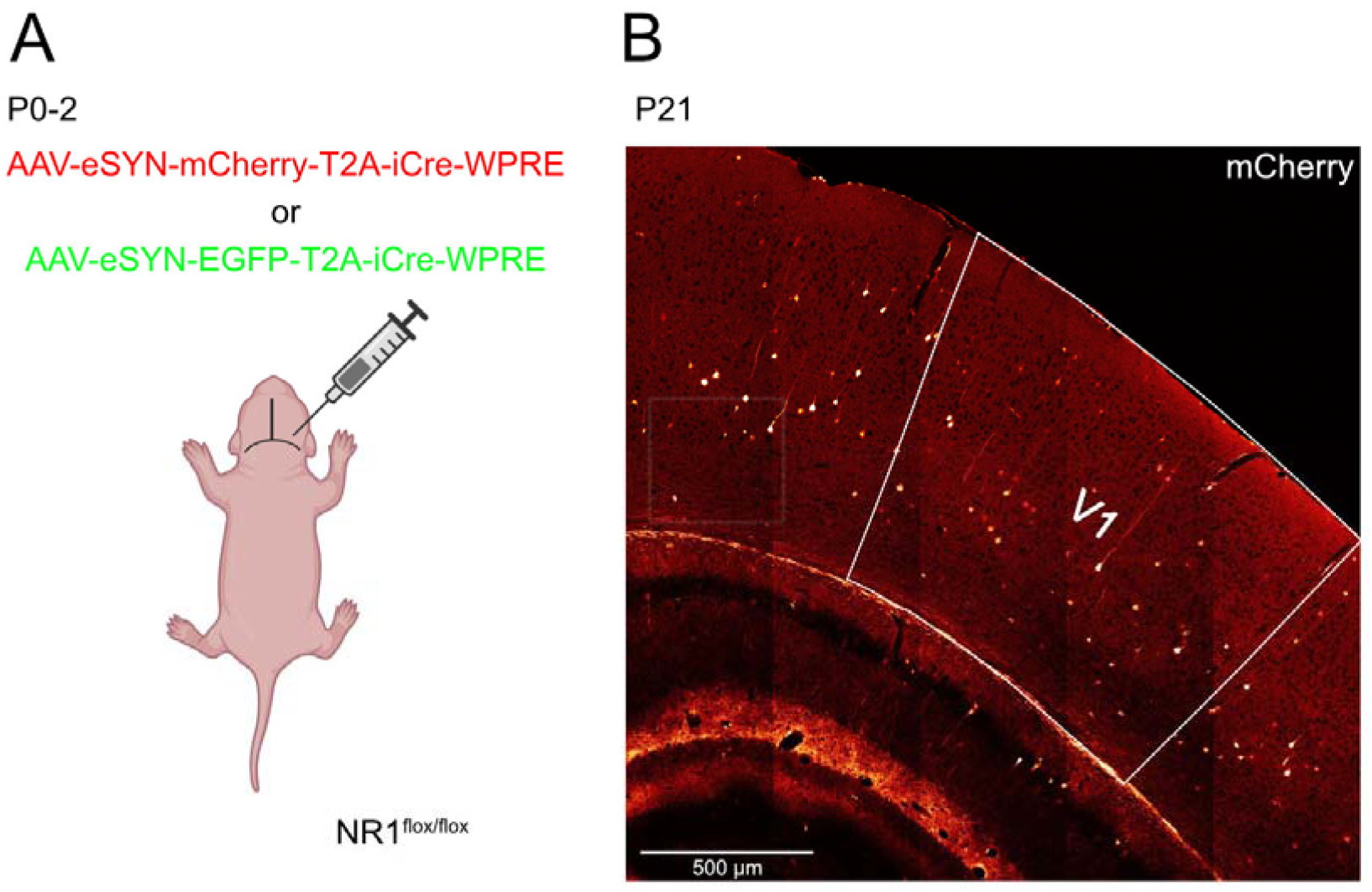
Sparse viral-mediated deletion of NMDARs in V1 excitatory neurons, Related to Figure 3. (A) Schematic of neonatal viral injection (P0–P2) delivering Cre recombinase and an mCherry reporter (AAV-eSYN-mCherry-T2A-iCre-WPRE) or EGFP reporter (AAV-eSYN-EGFP-T2A-iCre-WPRE) to excitatory neurons in primary visual cortex (V1) of NR1^flox/flox^ mice. Created in BioRender. Rannio, S. (2026) https://BioRender.com/27fmwdc (B) Representative confocal image showing sparse mCherry expression in V1 at P21, including labeling of a few L5 PCs with easily identified by their prominent apical dendrites.

**Supplementary Figure S6:**
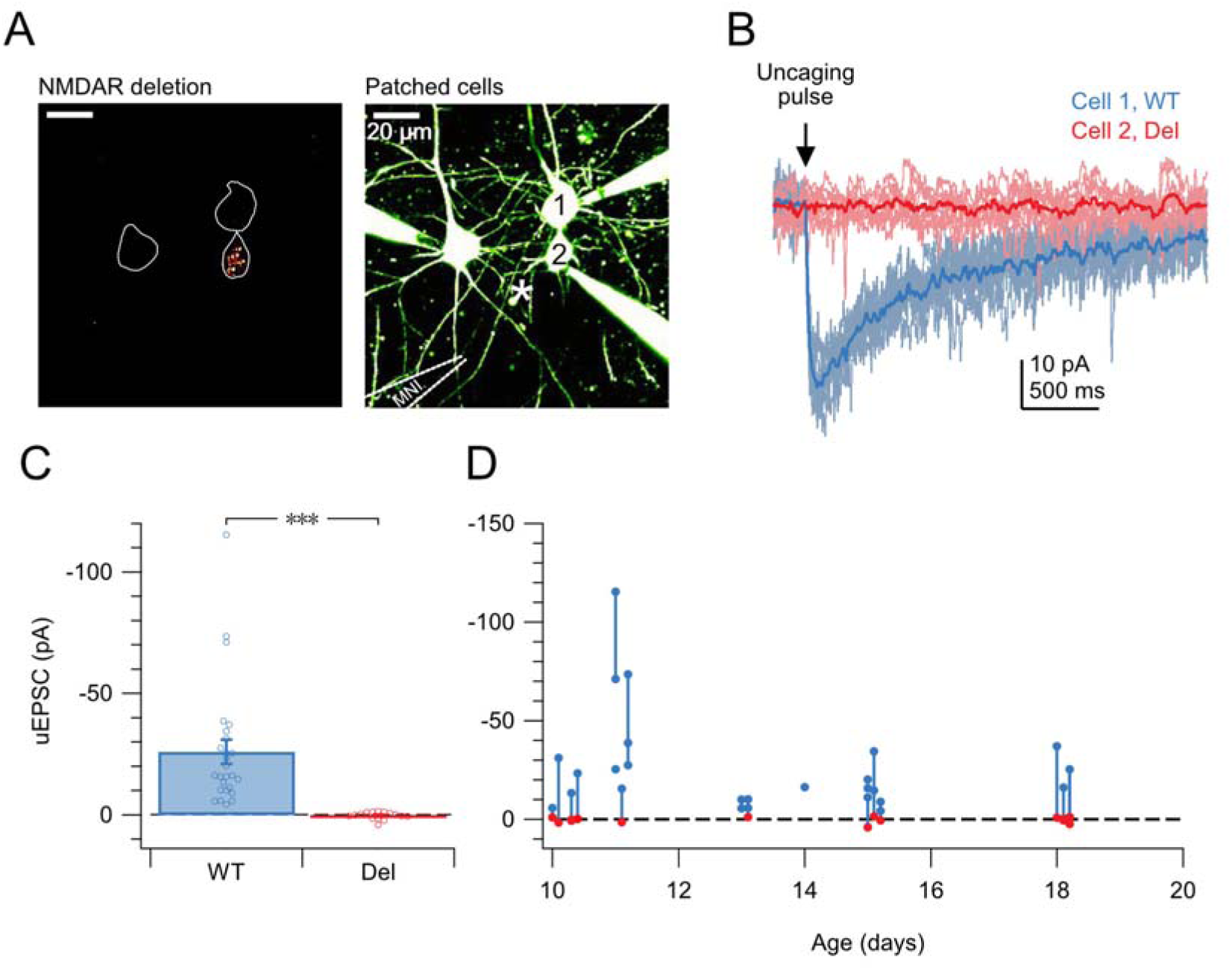
Uncaging confirmed efficient sparse GluN1 deletion, Related to Figure 3. (A) Left: mCherry fluorescence identifying PCs expressing AAV9-eSYN-mCherry-T2A-iCre-WPRE (see Methods). Cell outlines are based on Alexa 488 fills. Right: Two-photon image of V1 L5 PCs filled with Alexa Fluor 488, indicating the position of the laser uncaging site (*). MNI-NMDA was locally puffed near the uncaging location (“MNI”). (B) Ten-ms-long 405-nm laser pulses uncaged NMDA, resulting in NMDAR-mediated currents in PC1 but no detectable response in PC2 (averages of 10 sweeps shown), demonstrating that injection of AAV9-eSYN-mCherry-T2A-iCre-WPRE into NR1^fl/fl^ neonates (see Methods) deleted NMDARs in mCherry-tagged PCs. (C) Sparse viral delivery of Cre recombinase into NR1^flox/flox^ neonates reliably abolished uncaging-evoked NMDAR currents (uEPSCs) in V1 L5 PCs (*n*_WT_ = 26, *n*_Del_ = 15, Student’s t test). Each data point represents the mean of 10–30 sweeps. (D) From P10 onward, mCherry-positive PCs (red) showed no response to NMDA uncaging, whereas WT PCs (blue) responded robustly. Connected symbols indicate simultaneously recorded cell pairs. Each recording from a deleted PC was paired with a WT PC as a positive control for successful uncaging.

**Supplementary Figure S7:**
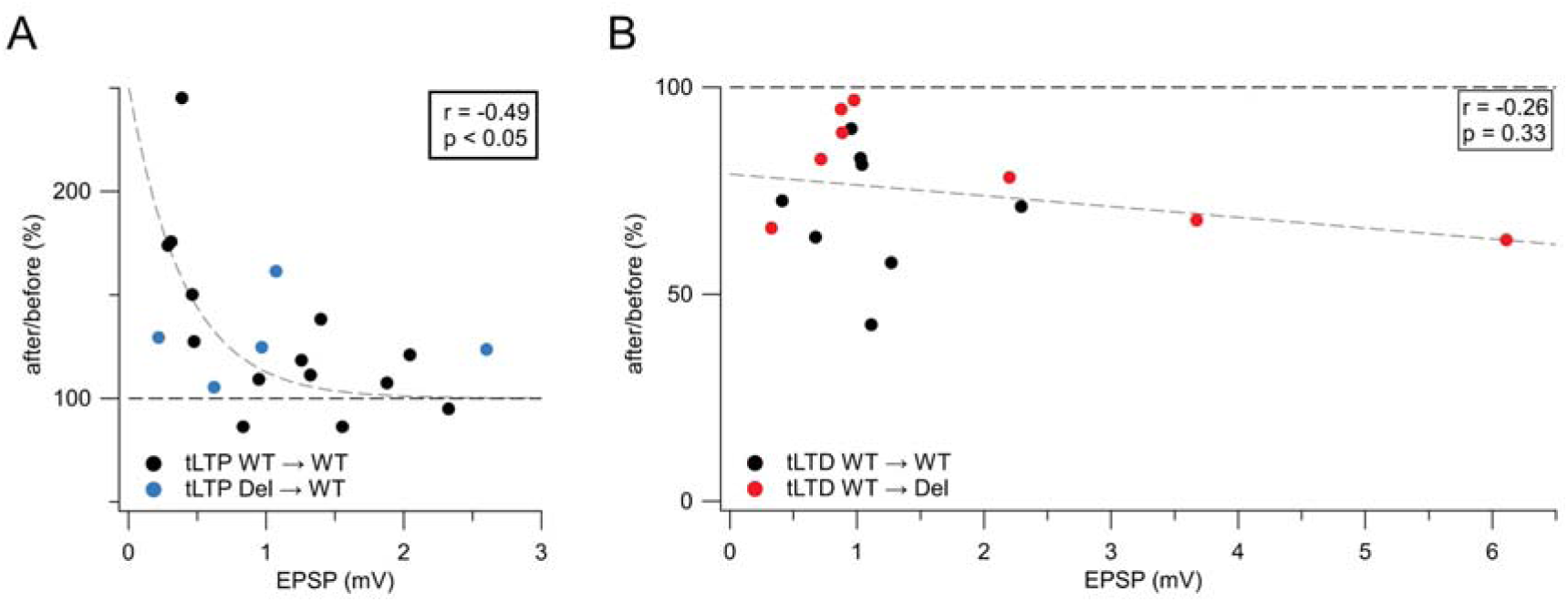
Initial synaptic strength predicted tLTP but not tLTD, Related to Figures 5-6. (A) As previously demonstrated for L5 PC → PC connections in the rat^10^ as well as for several other synapse types,^48–53^ weak connections potentiated more than strong ones (Student’s *t* for Pearson’s *r*), demonstrating saturation of potentiation for sufficiently strong synapses. (B) However, tLTD did not depend on synaptic strength, in agreement with previous findings.^10^ The properties of tLTP and tLTD thus agree with hallmark features in the prior literature.

**Supplementary Figure S8:**
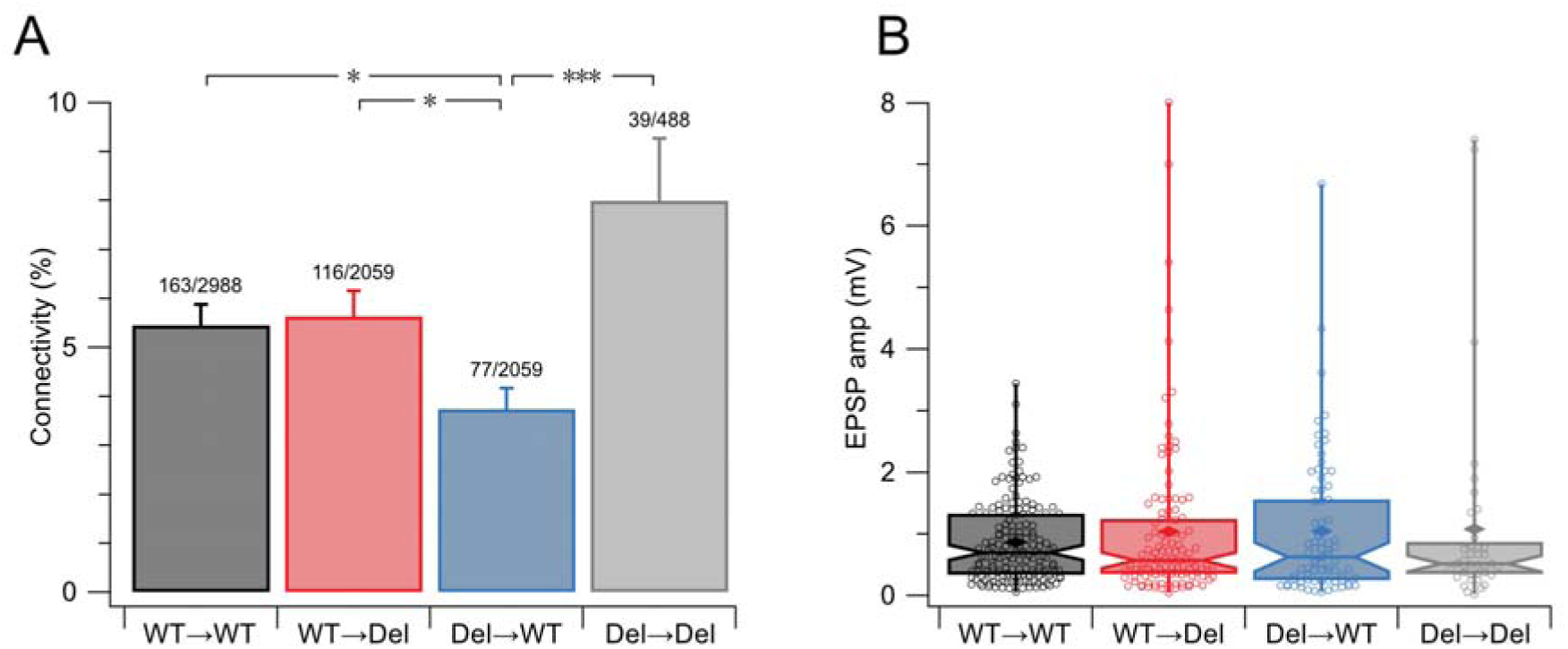
Connectivity rates, but not synaptic strength, differed with presynaptic NMDAR deletion in juvenile animals, Related to Figure 7. (A) WT → WT and WT → Del connectivity rates were indistinguishable (χ^2^ test, p = 0.78), indicating that postsynaptic NMDAR deletion did not measurably affect juvenile L5 PC → PC connectivity. By contrast, Del → WT connectivity differed from the other connection categories (Bonferroni-corrected χ^2^ tests), indicating a context-dependent role for presynaptic NMDARs in regulating connectivity. Error bars indicate normalized square root of counts. Absolute connectivity rates were lower here than we previously reported for L5 PC → PC pairs (10–17%),^10,38,57,93^ reflecting a deliberate experimenter bias toward superficially located neurons to enable reliable detection of fluorescently labeled cells, which likely increased axonal and dendritic truncation due to slicing. Relative comparisons across the four categories therefore remain interpretable. Reciprocal PC ↔L PC connections were examined but not quantified further due to low counts. (B) Sparse NMDAR deletion did not affect synaptic strength, as measured by EPSP amplitude (Kruskal-Wallis p = 0.903). Box plots show median and interquartile range, diamonds denote mean values, and whiskers indicate minimum and maximum values.

**Supplementary Figure S9:**
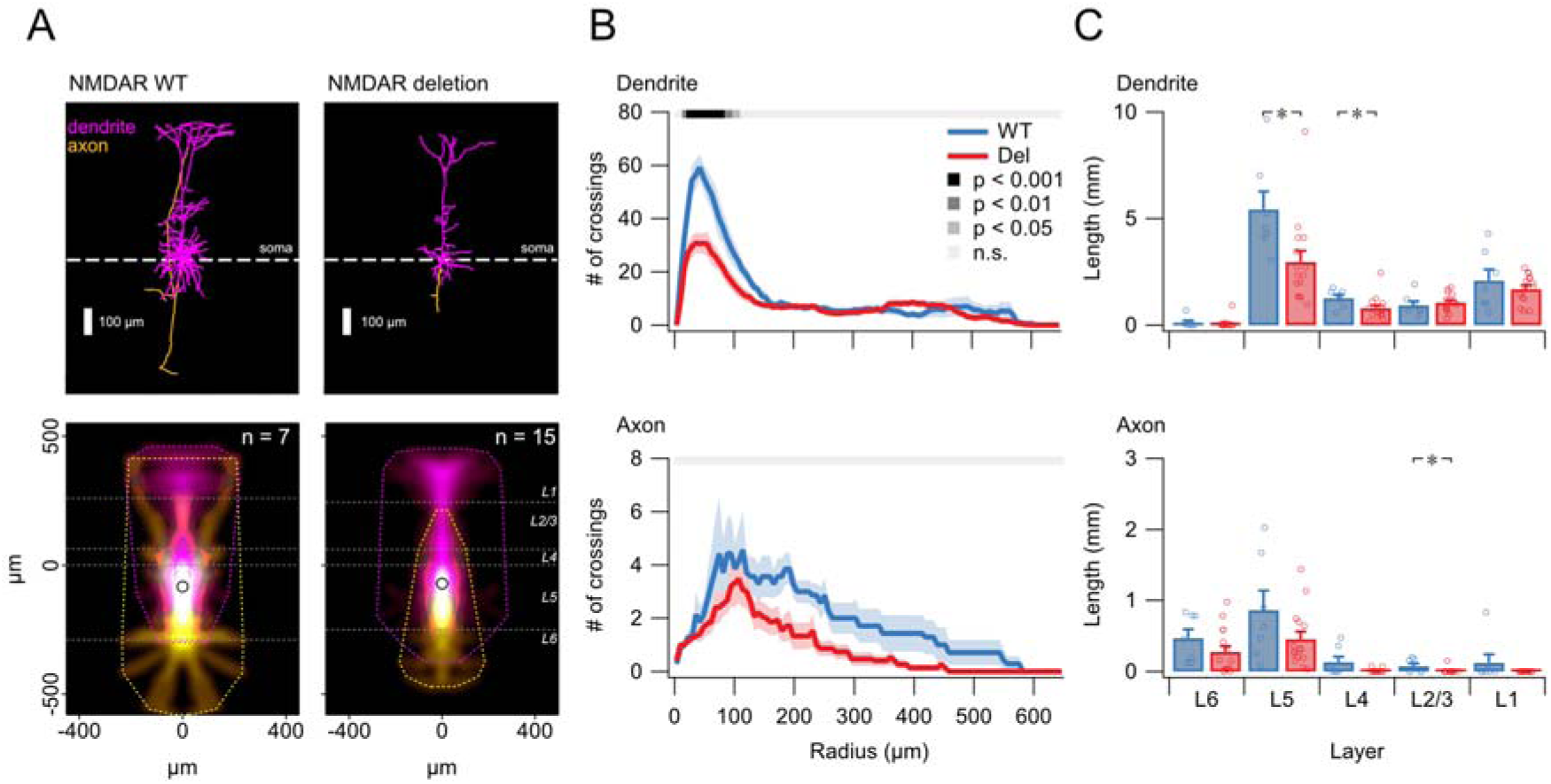
Global NMDAR deletion in juvenile mice recapitulates axonal and dendritic branching phenotypes, Related to Figure 8. (A) Sample morphologies of a WT PC (left) and a PC with global genetic NMDAR deletion. Reconstructions were aligned on their somata (dashed line). Bottom: Density maps show average extent of axonal (yellow) and dendritic (magenta) arborizations, while the convex hulls (yellow/magenta dotted lines) denote their maximum extents. Horizontal grey dotted lines demarcate neocortical layer boundaries. Open circles denote mean soma position. (B) LMM analysis of Sholl plot profiles demonstrated that NMDAR deletion specifically disrupted basal dendritic branching (top) but was not powered enough to conclusively reveal differences in axonal branching (bottom). The absence of a detectable axonal phenotype may reflect limited statistical power and challenges associated with tracing fine axonal processes. (C) Global genetic NMDAR reduced dendritic length in L4 and L5, as well as axonal length in L2/3 (one-tailed Student’s *t* test).

**Supplementary Figure S10:**
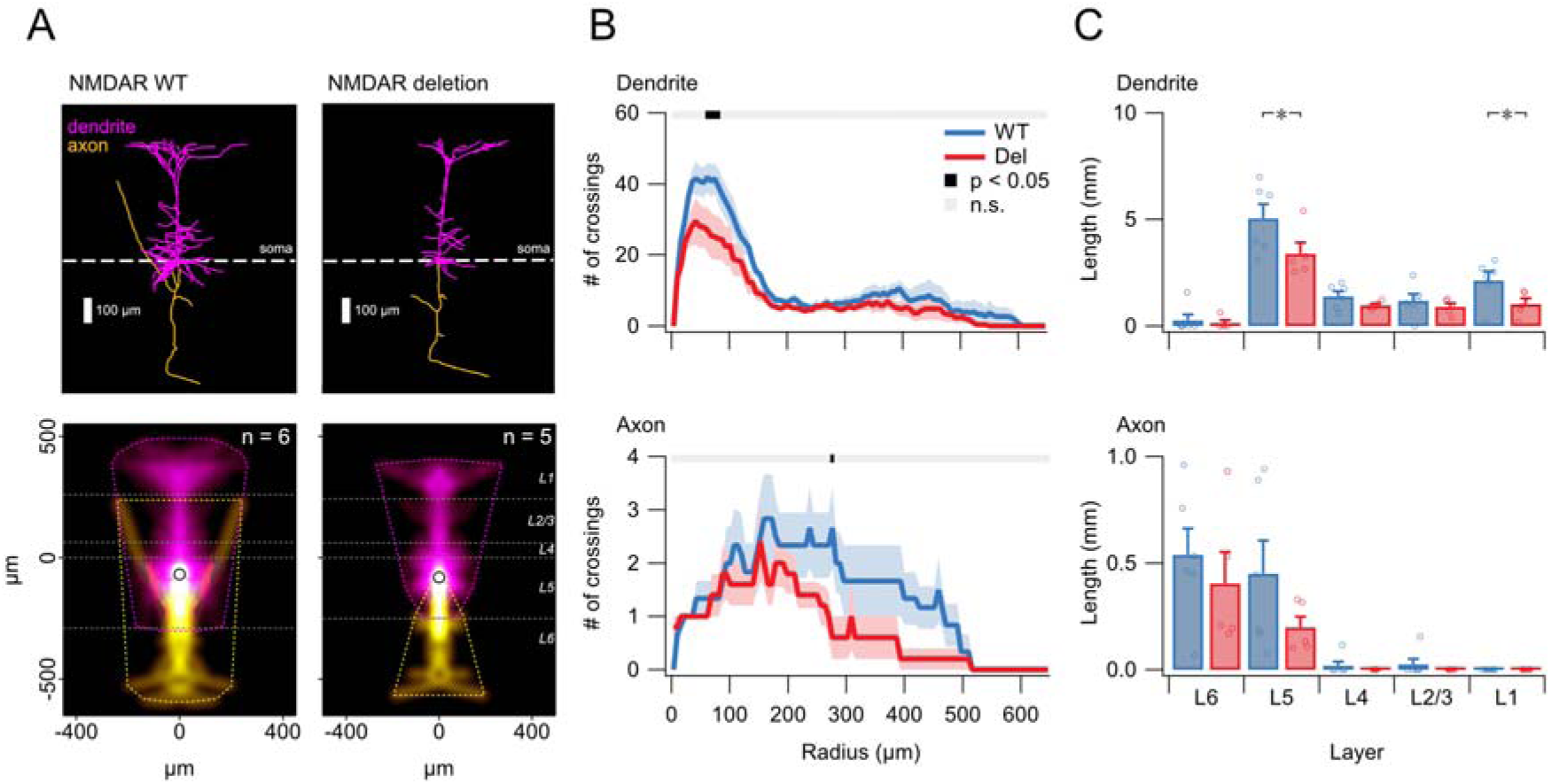
Sparse NMDAR deletion in adolescent mice supports layer-specific effects on axonal and dendritic branching, Related to Figure 8. (A) Sample morphologies of a WT PC (left) and a sparse genetic NMDAR deletion PC (right) in adolescent (P30-35) animals aligned on their somata. Bottom: Density maps show average extent of axonal (yellow) and dendritic (magenta) arborizations, while the convex hulls (yellow/magenta dotted lines) denote their maximum extents. Horizontal grey dotted lines demarcate neocortical layer boundaries. Open circles denote mean soma position. (B) LMM analysis of Sholl plot profiles suggested that sparse genetic NMDAR deletion disrupted branching in both dendrites (top) and axons (bottom). (C) Top: Sparse genetic NMDAR deletion reduced dendritic length in layers 1 and 5 (one-tailed t-test p < 0.05). Bottom: We did not observe a difference in axonal length across layers, although the absence of a detectable axonal phenotype may reflect limited statistical power and challenges associated with tracing fine axonal processes.

**Table S1:** NMDAR deletion did not affect spontaneous release rates in juvenile animals, Related to Figure 7.

| Category | Frequency (Hz) | Amplitude (pA) | Age (days) | *n*<sub>cells</sub> |
| --- | --- | --- | --- | --- |
| Global NMDAR deletion PC | 2.8 ± 0.1 | -10 ± 0.3 | 15 ± 0.2 | 86 |
| WT littermates PC | 2.7 ± 0.2 | -10 ± 0.3 | 15 ± 0.2 | 62 |
| Sparse NMDAR deletion PCs | 2.9 ± 0.3 | -10 ± 0.5 | 15 ± 0.3 | 38 |
| In-slice WT PCs | 2.9 ± 0.2 | -10 ± 0.4 | 16 ± 0.3 | 63 |

In juvenile L5 PC recordings from global NMDAR deletion mice (Emx1<sup>Cre/+</sup>;NR1<sup>fl/fl</sup>;Ai9<sup>tdTom/+</sup>) and WT littermates (Emx1<sup>Cre/+</sup>;NR1<sup>+/+</sup>;Ai9<sup>tdTom/+</sup>), the baseline mEPSC frequency as well as amplitude were indistinguishable (p = 0.40 and p = 0.66, respectively).
Likewise, sparse NMDAR deletion did not affect baseline mEPSC frequency (p = 0.62) or amplitude (p = 0.35), as assessed by comparing recombined PCs (NR1<sup>fl/fl</sup> + AAV9-eSYN-mCherry-T2A-iCre-WPRE) to in-slice nearby WT PCs (unrecombined NR1<sup>fl/fl</sup>).
Statistical comparisons were carried out with Mann-Whitney U tests for frequency and Student's t test for amplitude.

